# Comparative analysis between two GT4 glycosyltransferases related to polysaccharide biosynthesis in *Rhodococcus jostii* RHA1

**DOI:** 10.1101/2023.02.13.525685

**Authors:** Antonela E. Cereijo, María V. Ferretti, Alberto A. Iglesias, Héctor M. Álvarez, Matías D. Asencion Diez

## Abstract

The bacterial genus *Rhodococcus* comprises organisms that perform an oleaginous behavior under certain growth conditions and the ratio of carbon and nitrogen availability. Thus, *Rhodococcus* spp have outstanding biotechnological features as microbial producers of biofuel precursors, which would be used instead of lipids from crops. It was postulated that lipid and glycogen metabolism in *Rhodococci* are closely related. Thus, a better understanding of rhodococcal carbon partitioning requires identifying the catalytic steps redirecting sugar moieties to temporal storage molecules, such as glycogen and trehalose. In this work, we analyzed two glycosyl-transferases GT4 from *R. jostii*, *Rjo*GlgAb and *Rjo*GlgAc, which were annotated as α-glucan-α-1,4-glucosyl transferases, putatively involved in glycogen synthesis. Both enzymes were recombinantly produced in *E. coli* BL21 (DE3) cells, purified to near homogeneity, and kinetically characterized. *Rjo*GlgAb and *Rjo*GlgAc presented the “canonical” glycogen synthase (EC 2.4.1.21) activity. Besides, both enzymes were actives as maltose-1P synthases (GlgM, EC 2.4.1.342), although to a different extent. In this scenario, *Rjo*GlgAc is a homologous enzyme to the mycobacterial GlgM, with similar behavior regarding kinetic parameters and glucosyl-donor (ADP-glucose) preference. *Rjo*GlgAc was two orders of magnitude more efficient to glucosylate glucose-1P than glycogen. Also, this rhodococcal enzyme used glucosamine-1P as a catalytically efficient aglycon. On the other hand, both activities exhibited by *Rjo*GlgAb depicted similar kinetic efficiency and a preference for short-branched α-1,4-glucans. Curiously, *Rjo*GlgAb presented a super-oligomeric conformation (higher than 15 subunits), representing a novel enzyme with a unique structure to function relationships. Results presented herein constitute a milestone regarding polysaccharide biosynthesis in Actinobacteria, leading to (re)discovery of methyl-glucose lipo-polysaccharide metabolism in *Rhodococci*.

## Introduction

Gram-positive organisms with high G+C content in their genomes are significant constituents of the *phylum* Actinobacteria, including the *Rhodococcus* and *Mycobacterium* genera. It was described that several *Rhodococci* possess large genomes and non-chromosomal elements that, together, provide a wide potential for inhabiting different environments [1–3]. Individuals from *Rhodococcus* spp. were isolated from diverse niches, *e*.*g*., soil, water, and contaminated environments [4]. This notorious colonizing potential is usually ascribed to the extensive metabolic capacity presented by *Rhodococci*, which is related to their relatively large genomes and the presence of genes encoding unique catabolic and anabolic pathways [4, 5]. Besides, these organisms are known for their inherent ability to degrade organic compounds and aliphatic molecules [6, 7] or to synthesize “special” biomolecules, such as waxes and/or biolubricants [8, 9]. Thus, *Rhodococci* constitute a reference model regarding the design of bioremediation and/or microbial cell factory processes [10]. The increasing availability of molecular tools to modify rhodococcal genomes, and their metabolism, sustains the model role mentioned above [4, 11]. To improve such a task, it is critical to correlate the advance of genetic tools with a deeper knowledge of the intricated rhodococcal physiology. In addition, *Rhodoccoci* are phylogenetically nearby to the *Mycobacterium* genus. Then, the study of rhodococcal biochemistry could also be extrapolated to understand mycobacterial metabolic features, particularly regarding carbon directed to synthesize mycolic acids and the cell wall envelope [7,12,13].

The biochemical potential of *Rhodococci* coincides with multiple copies of genes for a certain function [5]. This *a priori* gene “redundancy” could be the base for rhodoccocal catabolic versatility, and some cases were associated with niche adaptation [5, 14]. In general, gene duplication may present functional redundancy and carry out the same biochemical feature in the cell. If the gene encodes an enzyme, the latter shows overlapping substrate ranges. Gene duplication can also evolve and encode similar enzymes, or isoenzymes, with different kinetics, substrates preference, and/or expression profiles [4]. These gene duplications are usually relatively recent or were under low evolutionary pressure. Then, gene duplication and specialization are at the basis of metabolism evolution and the appearance of new biocatalytic properties [15, 16]. Indeed, gene duplication and protein promiscuity are critical for offering a source of diversity and metabolic innovation in prokaryotes [17, 18].

The above-mentioned reinforces the importance of characterizing enzymes at the same or similar biochemical steps, as presented herein. Previous reports regarding the analysis of duplicated rhodococcal enzymes linked to carbohydrate metabolism exemplify that different kinetic behaviors arise after gene duplication [19, 20]. Tischler and collaborators [19] characterized the two trehalose-6P synthases (EC 2.4.1.15), OtsA1 and OtsA2, present in *Rhodococcus opacus*. The kinetic comparison showed that OtsA1 was highly active with UDP-glucose (UDP-Glc), as most bacterial OtsAs [21], while OtsA2 presented a preference for GDP-Glc as the substrate. In a recent work [20], our group showed that gene duplication at the level of UDP-Glc pyrophosphorylase (UDP-GlcPPase) is not redundant, and the study led to the discovery of a new type of enzyme specific for glucosamine-1P (GlcN-1P).

In rhodococcal cells, glycogen accumulation is linked to lipid and triacylglycerol metabolisms, acting as a temporal carbon allocation molecule [22, 23]. Two examples sustain the link mentioned above between both metabolisms: (*i*) intracellular glycogen highly accumulates after inhibition by cerulenin of fatty acids synthesis [22]; and (*ii*) NADPH, the reducing power fueling lipid biosynthesis is the primary inhibitor of rhodococcal ADP-GlcPPase, catalyzing the key step for glycogen synthesis [24]. On the other hand, methyl glucose lipopolysaccharide (MGLP) is a family of polysaccharides described in the *Mycobacterium* genus and some *Nocardia* [25, 26]. The structural basis of MGLPs is similar to glycogen, composed of α-1,4-linked glucose moieties (about 15 to 20 units), where some of those residues are 6-O-methylated and/or present different acylation degrees. It has been postulated that MGLP may regulate fatty acid elongation at the level of fatty acid synthase (FAS-1), given their ability to form complexes with long-chain fatty acids *in vitro* [26]. Knowledge of intracellular glucan biosynthesis at the molecular and enzymological level is pivotal to further understand the central feature of oleaginous *Rhodococcus* species -lipid and TAGs accumulation-which could be of biotechnological relevance in the biofuel industry [10]. Glycogen and MGLPs possess the potential to interact with fatty acid metabolism *in vitro*[26, 27], opening an opportunity to explore the link between carbohydrate and lipid metabolisms and a probable impact on TAGs production for biodiesel purposes [10]. In this work, we analyzed two glycosyl-transferases from *R. jostii*, *Rjo*GlgAb and *Rjo*GlgAc, annotated as α-glucan-α-1,4-glucosyl transferases or glycogen synthases. Each gene encoding *Rjo*GlgAb or *Rjo*GlgAc is located adjacent to another gene related to glycogen metabolism [5]: *glgAb* is together with the one encoding a putative α-1,4-glucan branching enzyme GlgB-type (EC 2.4.1.18) while *glgAc* to the *glgC* gene coding for ADP-GlcPPase (EC 2.7.7.27), the enzyme catalyzing the key step in the classical bacterial glycogen synthesis pathway [24,28–30]. We present here that both GlgA enzymes from *R. jostii* depict maltose-1P synthase and “classical” glycogen synthase activities, although to a different extent. The kinetic characterization shows that *Rjo*GlgAc is a homologous enzyme to the recently described and crystallized mycobacterial maltose-1P synthase GlgM (EC 2.4.1.342) [31, 32]. We deepened in its enzymatic properties by studying the ability to use alternative substrates, describing GlcN-1P as a catalytically efficient aglycon alternative to the canonical substrate Glc-1P. In addition, the GT4 *Rjo*GlgAb was obtained in a soluble and active form. The latter possesses a high sequence identity with the Rv3032 protein from *M. tuberculosis*, a so far recalcitrant enzyme ascribed to MGLP metabolism. We demonstrated its preference for short-branched α-1,4-glucans, in accordance to structural features proposed for actinobacterial glycogens. Curiously, *Rjo*GlgAb presented a super-oligomeric conformation with more than 15 subunits. This work shed light on the differential steps for synthesizing intracellular polysaccharides. Results are discussed regarding its impact on the comprehension of carbon partitioning in *R. jostii*, particularly carbohydrate metabolism related to lipid and/or TAGs production, that is, the prominent biotechnological feature of this group of oleaginous bacteria.

## Materials and Methods

### Chemicals

Protein standards, antibiotics, IPTG, Glc-1P, GlcN-1P, N-acetyl-glucosamine (GlcNAc-1P), galactosamine-1P (GalN-1P), mannose-1P (Man-1P), fructose-1P (Fru-1P), galactose-1P (Gal-1P), ADP-Glc, UDP-Glc and glycogen from rabbit liver were obtained from Sigma-Aldrich (Saint Louis, MO, USA). Genbiotech synthesized oligonucleotides. All other reagents were of the highest quality available. Actinobacterial glycogen (from *R. jostii* and *M. smegmatis*) was purified using a previously described alkali treatment [33–36]. Briefly, each pellet of cells was washed with ice-cold water, then resuspended in water and treated with KOH 30% (w/v) for 90 min at 100°C. After cooling and neutralizing with acetic acid, the polysaccharides were precipitated with ethanol 96% (v/v) at -20°C overnight. The suspensions were centrifuged at 20,000 xg for 30 min, and then polysaccharides were resuspended in water. The final concentration of the extracted glycogen was determined by its digestion with amyloglucosidase (Sigma-Aldrich), and the released glucose was measured with the glucose oxidase method [37] using a commercial kit (Wiener Lab).

### Bacteria and plasmids

*Escherichia coli* Top 10 F’ cells (Invitrogen) and pGEM^®^-T Easy vector (Promega) were used for cloning procedures. The *glgAb* and *glgAc* genes from *R. jostii* were expressed in *Escherichia coli* BL21 (DE3) (Invitrogen) using a pET28c vector (Novagen). DNA manipulations, *E. coli* cultures, and transformations were performed according to standard protocols (Gehring et al., 1990).

### Gene amplification

The *glgAb* (ID 4224010) and *glgAc* (ID 4223525) genes coding for *R. jostii* RHA1 *Rjo*GlgAb and *Rjo*GlgAc, respectively, were amplified by PCR using genomic DNA as a template. Primers (detailed in Table S1) were designed according to available genomic information for *R. jostii*[5] in the GenBank database (http://www.ncbi.nlm.nih.gov/nuccore/111017022www.ncbi.nlm.nih.gov/Genbank/index.html). PCR reaction mixtures (50 µl) contained 100 ng of genomic DNA, 50 pmol of each primer, 0.2 mM of each dNTP, 2.5 mM Mg^2+^, 8% (v/v) DMSO, and 1 U *Taq* DNA polymerase (Fermentas). Standard conditions of PCR were used for 30 cycles: denaturation at 94 °C for 1 min, annealing for 42 s at 58.4 °C for both genes, and extension at 72 °C for 1.5 min, with a final extension of 10 min at 72 °C. PCR reaction mixtures were solved in 1% (w/v) agarose gel and purified using Wizard SV gel & PCR Clean-Up kits (Promega). Then, the amplified genes were cloned into the T-tailed plasmid pGEM-T Easy and their identities were determined by DNA sequencing (Macrogen, Korea).

### Cloning, expression, and purification procedures

The pGEM-T Easy plasmids harboring either *glgAb* or *glgAc* from *R. jostii* were digested with the corresponding restriction enzyme (see Table S1) and cloned into pET28c vector to obtain the expression constructions [pET28c/*RjoglgAb*] and [pET28c/*RjoglgAc*]. Competent *E. coli* BL21 (DE3) cells were then transformed with single constructions for individual *Rjo*GlgAb or *Rjo*GlgAc production. Protein expression was carried out using LB medium (10 g/l tryptone; 5 g/l yeast extract; 10 g/l NaCl) supplemented with 50 µg/ml kanamycin. Cells were grown at 37 °C and 250 rpm until OD_600_ ∼0.6. Recombinant gene expression was induced for 16 h at 20 °C by adding 0.2 mM IPTG. After induction, cells were harvested by centrifugation at 5,000 *×g* for 10 min and stored at -20 °C until use. His-tagged proteins were purified by immobilized-metal ion affinity chromatography (IMAC), where all purification steps were performed at 4 °C. After resuspension in buffer H (50 mM Tris-HCl [pH 8.0], 300 mM NaCl, 10 mM imidazole, 5% [vol/vol] glycerol), cells were disrupted by sonication (5-s pulse on with intervals of 3-s pulse off for a total time of 10 min on ice) and later centrifuged twice (10 min) at 30,000 *×g*. Supernatants were loaded in a 1-ml His-Trap column (GE Healthcare) previously equilibrated with buffer H. The recombinant proteins were eluted with a 10 to 200 mM imidazole linear gradient in buffer H (50 volumes). Fractions containing the highest activity were pooled, concentrated, and dialyzed against buffer H. The resulting enzyme samples were stored at -80 °C until use, remaining fully active for at least six months.

### Molecular mass determination

Protein molecular mass at the native state was determined by gel filtration using a Superdex200 10/300 column (GE Healthcare). A gel filtration calibration kit (high molecular weight; GE Healthcare) with protein standards including thyroglobulin (669 kDa), ferritin (440 kDa), aldolase (158 kDa), conalbumin (75 kDa), and ovalbumin (44 kDa) was used. The column’s void volume was determined using Dextran Blue (Promega).

### Protein measurement

Protein concentration was determined by the modified Bradford assay [38] using BSA as a standard. Recombinant proteins and purification fractions were defined by sodium dodecyl sulfate-polyacrylamide gel electrophoresis (SDS-PAGE), according to [39]. Gels were loaded with 5 to 50 µg of protein per well and stained with Coomassie-Brilliant Blue.

### Enzyme activity assays

The activity was determined at 37 °C using a continuous method where the NDP formation is enzymatically coupled to NADH consumption, as previously described for other glycosyl-transferases [40–42]. Pyruvate kinase from rabbit muscle (PK, Sigma) and lactate dehydrogenase from *Lactobacillus* (LDH, Sigma) were used as coupling enzymes. The reaction mixture contained 50 mM MOPS pH 8.0, 10 mM MgCl_2_, 0.3 mM PEP and 0.3 mM NADH; ADP-Glc, 0.02 U/µl PK, 0.02 U/µl, and a proper enzyme dilution. With a total volume of 50 µl, assays were initiated by adding glycogen from rabbit liver or by the corresponding sugar-1P. All the enzymatic assays were performed for 10 min at 37 °C in a 384-well multiwell plate (NuncTM), measuring the absorbance at 340 nm with a spectrophotometer Multiskan GO (Thermo Scientific).

One unit of activity (U) is defined as the amount of enzyme catalyzing the formation of 1 µmol of product per min under the conditions above described in each case.

### Calculation of kinetic constants

Saturation curves were performed by assaying enzyme activity at different concentrations of the variable substrate or effector and saturating levels of the others. Experimental data were plotted as enzyme activity (U/mg) *versus* substrate concentration (mM), and kinetic constants were determined by fitting the data to the Hill equation as described elsewhere [43]. Fitting was performed with the Levenberg-Marquardt nonlinear least-squares algorithm provided by the computer program Origin^TM^. Hill plots were used to calculate the Hill coefficient (*n*_H_), the maximal velocity (*V*_max_), and the kinetic constants that belong to substrate concentrations giving 50% of the maximal velocity (*K*_m_). The *k*_cat_ values were calculated considering a single catalytic subunit for each enzyme (44.86 kDa and 41.37 kDa for *Rjo*GlgAc and *Rjo*GlgAb, respectively). All kinetic constants are the mean of at least three independent data sets of reproducible within ±10%.

### Phylogenetic analysis

Amino acid sequences of different GlgA polypeptides from other organisms were downloaded from the NCBI database (http://www.ncbi.nlm.nih.gov). They were filtered to remove duplicates and near duplicates (*i.e*., mutants and strains from the same species). A preliminary alignment was constructed using the ClustalW multiple-sequence alignment server [44]. Sequences with an incorrect annotation or that were truncated were also eliminated manually. After this, sequences were manually refined with the BioEdit 7.0 program [45]. The tree was constructed using SeaView 4 [46] with the neighbor-joining algorithm (bootstrap of 1000). Confidence coefficients for the tree branches were obtained and plotted. Finally, the tree was prepared with the FigTree 1.3 program (http://tree.bio.ed.ac.uk).

## Results

### Theoretical analysis of the glgA duplication in R. jostii

*R. jostii* displays several gene duplications along its genome, some related to carbohydrate metabolism [5]. We recently demonstrated that the *galU* duplication at the level of hexoses-1P is not trivial since we reported an enzyme with a novel substrate specificity (GalU2), different from the canonical UDP-GlcPPase (EC 2.7.7.9) [20]. In this work, we extended the analysis to other metabolic steps from rhodococcal carbohydrate metabolism, such as the case of polysaccharide(s) synthesis. *R. jostii* possesses genetic fragments encoding putative glycogen synthases: *RjoglgAb* (ID 4224010) and *RjoglgAc* (ID 4223525) [5]. The predicted *Rjo*GlgAb and *Rjo*GlgAc proteins have theoretical molecular masses of 44.86 kDa and 41.37 kDa, respectively, with a 27% identity. *RjoglgAc* encodes a homolog of the mycobacterial GlgA with a 66% identity, already characterized [31, 41] and recently crystallized and renamed as GlgM [32]. *Rjo*GlgAb, instead, shows 33% identity with *M. tuberculosis* GlgM and 71% identity with the *M. tuberculosis* Rv3032 protein, which remains enzymatically uncharacterized so far. *Rjo*GlgAc shares 67.53% and 29.62% identities with GlgM and Rv3032 proteins from *M. tuberculosis*, respectively. On the other hand, *Rjo*GlgAc shows low identity values (21-24%) with the already characterized and crystallized glycogen synthases from *E. coli* and *A. tumefaciens*, while *Rjo*GlgAb has 26-28% identity towards the same enzymes [47–51]. Also, each rhodococcal GlgA studied herein shares about 25% identity with GlgA1 or GlgA2 from *Synechocystis*, a cyanobacterium reported to have duplicated glycogen synthases with differential biochemical behavior [52, 53]. Then, the comparative study in this work provides a milestone regarding glycosyl transferase activities related to (actino)bacterial polysaccharide synthesis.

We constructed an alignment (Supplementary Figure 1) with protein sequences from both rhodococcal GlgAs and those from solved structures, including GT4 mycobacterial GlgMs [32] and GT5 GlgAs from *E. coli* [50] and *Agrobacterium tumefaciens* [47]. We added to the analysis the GlgA1 and GlgA2 sequences from cyanobacterium with a duplicated glycogen synthase case [53, 54]. Given the GT-B fold they adopt, a structural similarity was established between the GT5 family of bacterial glycogen synthases and mycobacterial GT4 GlgMs, since both types of enzymes share the glucosyl donor ADP-Glc [32, 55]. The alignment shows that critical binding and catalytic amino acid residues described in GlgA and GlgM proteins are conserved either in *Rjo*GlgAc or *Rjo*GlgAb. Supplementary Figure 1 also illustrates that *Rjo*GlgAc harbors the same amino acids proposed to interact with ADP-Glc in the crystallized GlgM. Instead, *Rjo*GlgAb possesses all the identical conserved residues but only a Val to Cys (Cys161) mutation (Val146 in the *M. smegmatis* enzyme). The “Val146” is also mutated to a Cys residue in the mycobacterial Rv3032, thus reinforcing the similitude between both mycobacterial and rhodococcal proteins. As a difference, Rv3032 has a Leu instead of the Ile293 of the mycobacterial GlgM. Curiously, this Leu residue is also present in the glycogen synthases GT5 studied (Supplementary Figure 1). These sequence similitudes amongst the analyzed glycosyl-transferases sustain the importance of advancing the study of structure-to-function relationships in the actinobacterial GlgA-type enzymes. The arising question refers to how these proteins diverged in their ability to elongate/produce (actino)bacterial glycans.

### Recombinant protein production, purification, and molecular mass determination

The *RjoglgAb* (1,245 bp) and *RjoglgAc* (1,170 bp) genes were amplified from *R. jostii* genomic DNA with individual single-step PCR procedures, and their identities were confirmed by DNA sequencing. After cloning each gene into the pET28 plasmid, we obtained suitable vectors for the independent heterologous protein production in *E. coli*. The recombinant expression in *E. coli* BL21 (DE3) showed the production of both *Rjo*GlgAb and *Rjo*GlgAc proteins in soluble fractions (not shown). Afterward, the purification strategies allowed the recovery of the enzymes with a high purity degree, as presented in Figure 1.A. During the purification steps, activity was followed by the “canonical” glucan elongation ability, confirming that both rhodococcal GlgAs were active as glycogen synthases. The *Rjo*GlgAb soluble production remarkably triggers advancing its structure-to-function characterization. Particularly, results obtained with this rhodococcal enzyme could be extrapolated to its homolog Rv3032 from *M. tuberculosis*, an uncharacterized enzyme associated with MGLP metabolism [26, 56].

**Figure 1:**
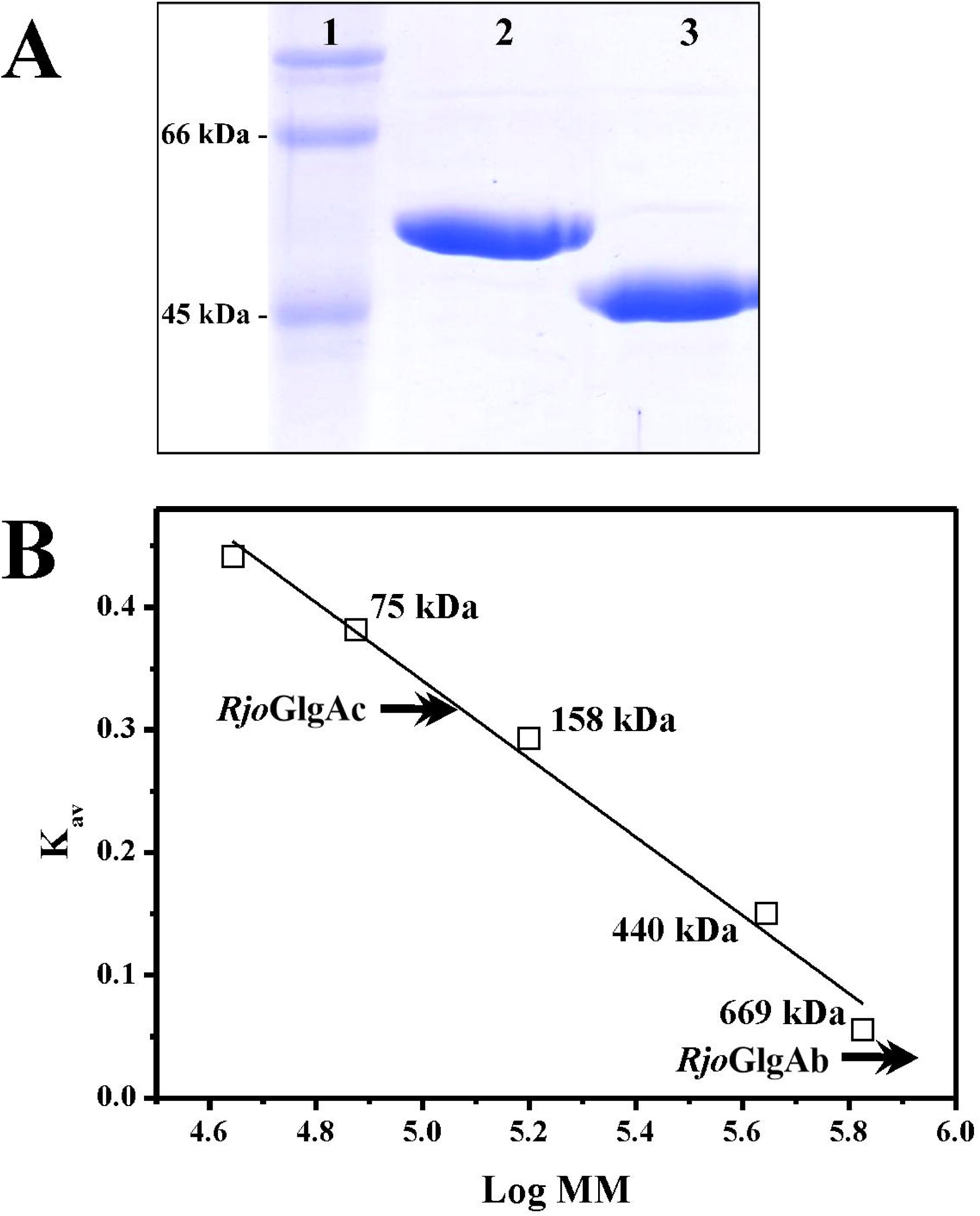
Purified rhodococcal GlgAs proteins analysis. **A:** SDS-PAGE (12%) of *Rjo*GlgAc **(lane 2)** and *Rjo*GlgAb **(lane 3)**; molecular mass markers **(lane 1)**. **B:** Molecular mass (MM) determination by size exclusion chromatography on a Superdex 200 column as detailed in Materials and Methods.

The purified GlgAs from *R. jostii* were analyzed by gel filtration to approach their quaternary structure determination. As shown in Figure 1.B, *Rjo*GlgAc eluted as a dimer, agreeing with previous results with the mycobacterial GlgM [32] and the GlgA from *S. coelicolor* [57]. On the other hand, *Rjo*GlgAb eluted at a volume lower than the largest marker (667 kDa) used during column calibration (Supplementary Figure 2). Considering the theoretical molecular mass of the *Rjo*GlgAb monomer (∼45 kDa), an oligomeric conformation of at least 15 subunits could be inferred. We confirmed that the eluted protein at a high oligomeric structure was the active conformation (Supplementary Figure 2). Then, further questions arise regarding the *Rjo*GlgAb structure-to-function relationships, which are aimed at future work. As presented here, the obtention of soluble enzymes is a remarkable output *per se*, offering molecular tools to advance in the comparative kinetic and structural characterization of two key proteins regarding polyglucan synthesis in Actinobacteria.

### Kinetic characterization: glycogen elongation

Since both rhodococcal GlgAs were annotated as putative glycogen synthases (the activity used to follow purification processes), *Rjo*GlgAc and *Rjo*GlgAb were analyzed regarding their “canonical” activity related to elongating an α-1,4-glucan chain. We confirmed that both enzymes were active as glycogen synthases (EC 2.4.1.21). Activity values of 0.25 U/mg and 1.1 U/mg for *Rjo*GlgAb and *Rjo*GlgAc were respectively obtained (at 1 mM ADP-Glc and 2 mg/ml of rabbit muscle glycogen in the reaction mixture). Then, both rhodococcal GlgA enzymes were characterized in the α-1,4-glucan elongation direction, with results detailed in Table 1. *Rjo*GlgAc is 3-fold more active, and its affinity toward glycogen is one order of magnitude higher than *Rjo*GlgAb. Also, *Rjo*GlgAc depicted a 6-fold higher apparent affinity for ADP-Glc than *Rjo*GlgAb (Table 1). Both rhodococcal GlgAs were highly specific for ADP-Glc since no activity was detected when up to 10 mM UDP-Glc or GDP-Glc were present in reaction mixtures containing 2 mg/ml of rabbit muscle glycogen.

**Table 1:**
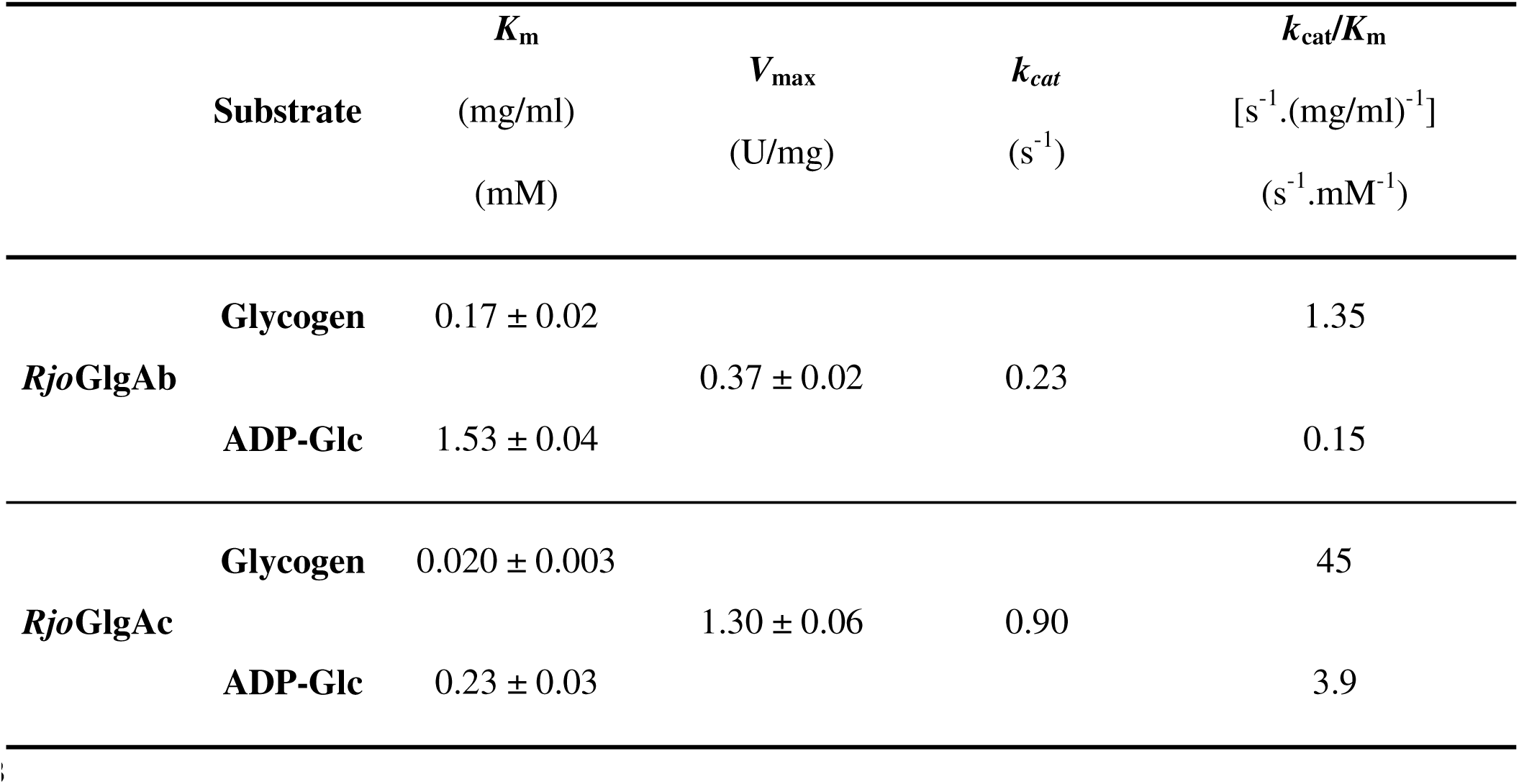
Kinetic parameters of both glycogen synthases from R. jostii, RjoGlgAb and RjoGlgAc. The K_m_ value is expressed in mg/ml or mM for glycogen and ADP-Glc, respectively. k_cat_ were calculated using the corresponding theoretical molecular mass of the monomer.

Given that *Rjo*GlgAc presented a high identity to mycobacterial GlgM (see below) and that *Rjo*GlgAb was an already uncharacterized enzyme, we emphasized the study of *Rjo*GlgAb kinetic properties analyzing different glucan molecules to broaden the knowledge regarding its polysaccharide preference. First, we analyzed maltose (the minimum α-1,4-glucan), cellobiose (β-1,4-bond), and glycogen from oyster (type III). The activity with maltose and cellobiose was either neglectable (about 50 mU/mg) or undetectable, respectively. Curiously, *Rjo*GlgAb displayed 2-fold higher activity when glycogen from oyster was assayed as an aglycon substrate, as shown in Supplementary Figure 3. Also, the *Rjo*GlgAb affinity for oyster glycogen increased ∼2-fold compared to the “control” with the one from rabbit muscle (see Figure 2 and Table 1). It was described that glycogen from oyster possesses a heterogeneous structure given the presence of glucan chains with different lengths, where seven glucose moieties are considered the average size [58, 59]. In contrast, glycogen from bacteria and eukaryotes is composed of identical branches with 12-14 glucose units [60–62].

**Figure 2:**
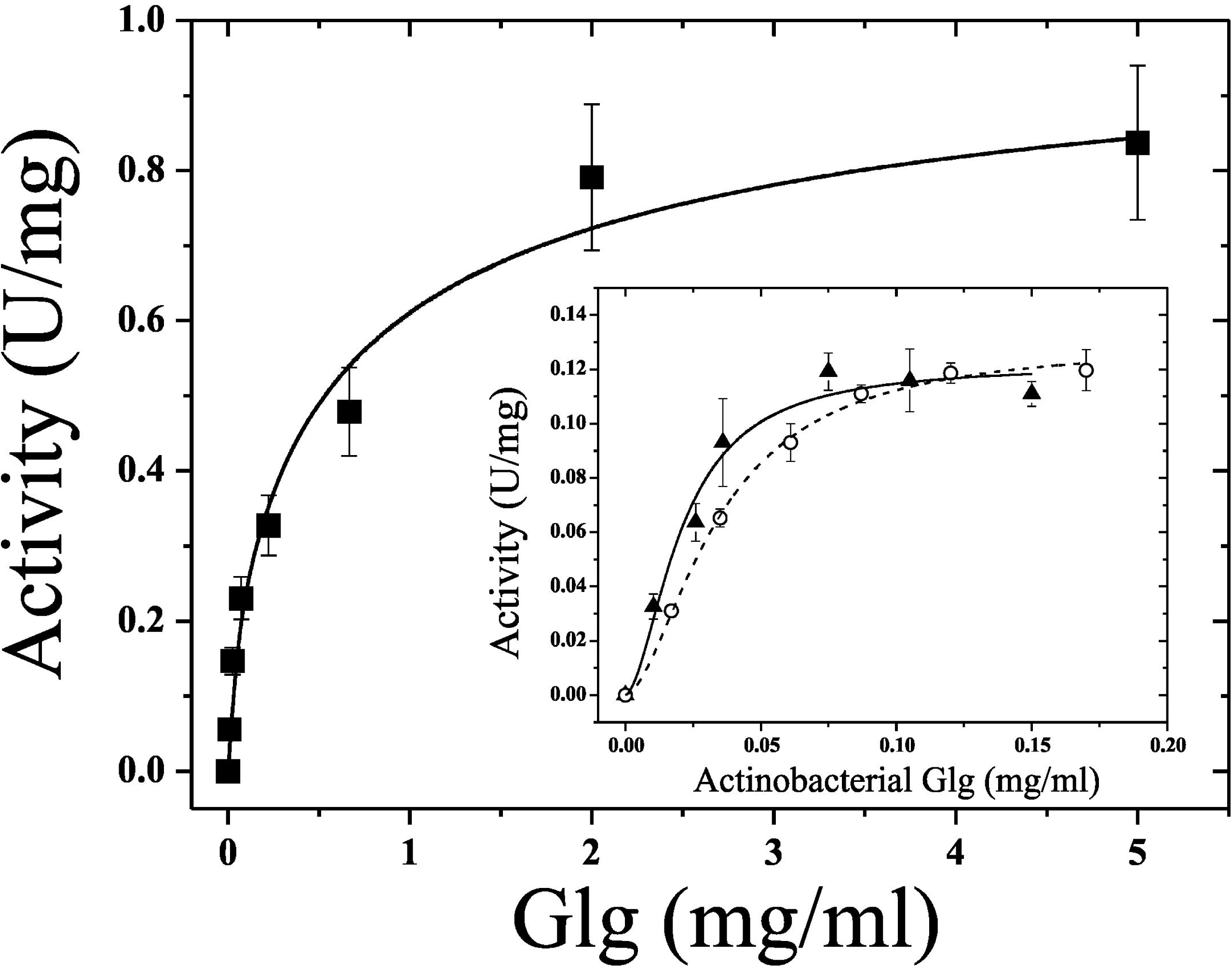
*Rjo*GlgAb saturation curves with different types of glycogens. Oyster glycogen saturation curve is presented in the figure, while the insert shows saturation curves made with glycogen purified from *R. jostii* (fill triangles, solid line) and from *M. smegmatis* (open circles, dash line). Curves were conducted using 5 mM ADP-Glc.

Worthy to note, the structural analysis of the intracellular glycogen from *M. tuberculosis* showed it is conformed with “short” α-1,6-glucosides branches, containing mainly between two and six residues [63]. Thus, we assayed polysaccharides extracted from Actinobacteria (*M. smegmatis* and *R. jostii*). The latter was incorporated, given that *Rjo*GlgAb (and Rv3032) analogs are only found in some actinobacterial members (see Discussion). The kinetic characterization with the replacement of rabbit muscle glycogen for those from an actinobacterial source showed no significant differences in activity. On the other hand, 5-to 8-fold higher apparent affinities for glycogens from bacterial sources were determined, with *K*_m_ values of 0.021 mg/ml and 0.034 mg/ml for the polysaccharides extracted from *R. jostii* and *M. smegmatis*, respectively, as shown in the inset of Figure 2. Then, *Rjo*GlgAb glucan preference indicates that shorter glucan branches elicit higher activity values and that specific structures present in actinobacterial glucans cause increased affinities.

Overall, results confirm the relatively low catalytic ability as canonical glycogen synthases to elongate glucan molecules from ADP-Glc, thus suggesting the appearance of an alternative specialized function, as presented below for *Rjo*GlgAc. The poor efficiency with glycogen molecules sustains the hypothesis that the *Rjo*GlgAb substrates might be related to MGLP metabolism, as proposed for the mycobacterial protein Rv3032 [26, 56]. While the parameters for *Rjo*GlgAb are closer to the values obtained in the characterization of the GlgA from *M. tuberculosis* [41], the measured *K*_m_ for *Rjo*GlgAc are similar to those obtained for the enzyme from *S. coelicolor* [57], being one of the highest affinities towards the polyglucan reported so far (see Table 1 and Supplementary Figure 4). Remarkably, *Rjo*GlgAb showed these high-affinity values when actinobacterial glycogens were used as substrates. In addition, *Rjo*GlgAc depicts a catalytic efficiency for glycogen elongation from ADP-Glc agreeing with metabolic feasibility [64, 65]. Taken together, results suggest the possibility of an active classic pathway (GlgC/GlgA) for glycogen synthesis in *R. jostii* [22], although in mycobacteria, little (if any) glycogen is produced by this pathway [31, 66]. *Rjo*GlgAb presents 71% identity with the mycobacterial Rv3032 protein, an α-1,4-glycosyltransferase from *M. tuberculosis* putatively involved in MGLP elongation [56,63,67]. Until now, there was no enzymatic data regarding Rv3032, although it has been suggested that the enzyme could use both ADP-Glc and UDP-Glc [63, 68].

### Kinetic characterization II: maltose-1P synthesis

It has been described the ability of the mycobacterial GlgA to synthesize maltose-1P using ADP-Glc and Glc-1P as substrates [31], leading to a new activity named maltose-1P synthase (GlgM; EC 2.4.1.342) [32]. Maltose-1P is the glycosyl donor for glycogen elongation via the maltosyl transferase GlgE (EC 2.4.99.16) [69, 70]. Until now, reported kinetic parameters for GlgM enzymes belong to mycobacterial sources, and no data is available with those from alternative microorganisms. In the case of *Rhodococci*, glycogen synthesis is interconnected with lipid production, which represents their potential as biofactories [10]. Then, we analyzed the rhodococcal GlgA enzymes’ ability to catalyze maltose-1P synthesis, thus incorporating new elements into carbon partitioning analysis in *R. jostii*.

As expected, given its 66% identity to mycobacterial GlgM, *Rjo*GlgAc was active as a maltose-1P synthase. Curiously, *Rjo*GlgAb also catalyzed the disaccharide-1P formation, although with a *k*_cat_ 500-fold lower. Despite, both enzymes depicted similar kinetic behavior (see Supplementary Figure 4) and parameters are presented in Table 2. *Rjo*GlgAc portrayed substrate inhibition for Glc-1P curves (Figure 3), in agreement with the mycobacterial GlgM enzymes [31, 32]. In the presence of 3 mM ADP-Glc, we observed a peak (∼150 U/mg) at 0.5 mM Glc-1P, which decreased 35% and 50% in activity at 1 mM and 2 mM, respectively. When the Glc-1P curve was assayed with 0.3 mM ADP-Glc (a concentration close to its *K*_m_ value, Table 2), a similar performance was detected (∼70 U/mg) where the activity then diminished 15% and 48% at 1mM and 2 mM Glc-1P, respectively (Figure 3A). *Rjo*GlgAb presented the inhibition pattern described for *Rjo*GlgAc when Glc-1P curves were analyzed, as shown in Figure 3B. With 1 mM ADP-Glc, an activity peak (0.27 U/mg) was present between 0.5-0.6 mM Glc-1P, which decreased 16% and 32% at concentrations belonging to 1 mM and 2 mM of the hexose-1P. The same approach, with 0.2 mM ADP-Glc, showed a 50% and 81% decrease at 1 and 2 mM Glc-1P, respectively. Then, although with different catalytic magnitudes, both GlgAs from *R. jostii* share the ability to produce maltose-1P with a common kinetic behavior, similar to previous reports [31, 32].

**Figure 3:**
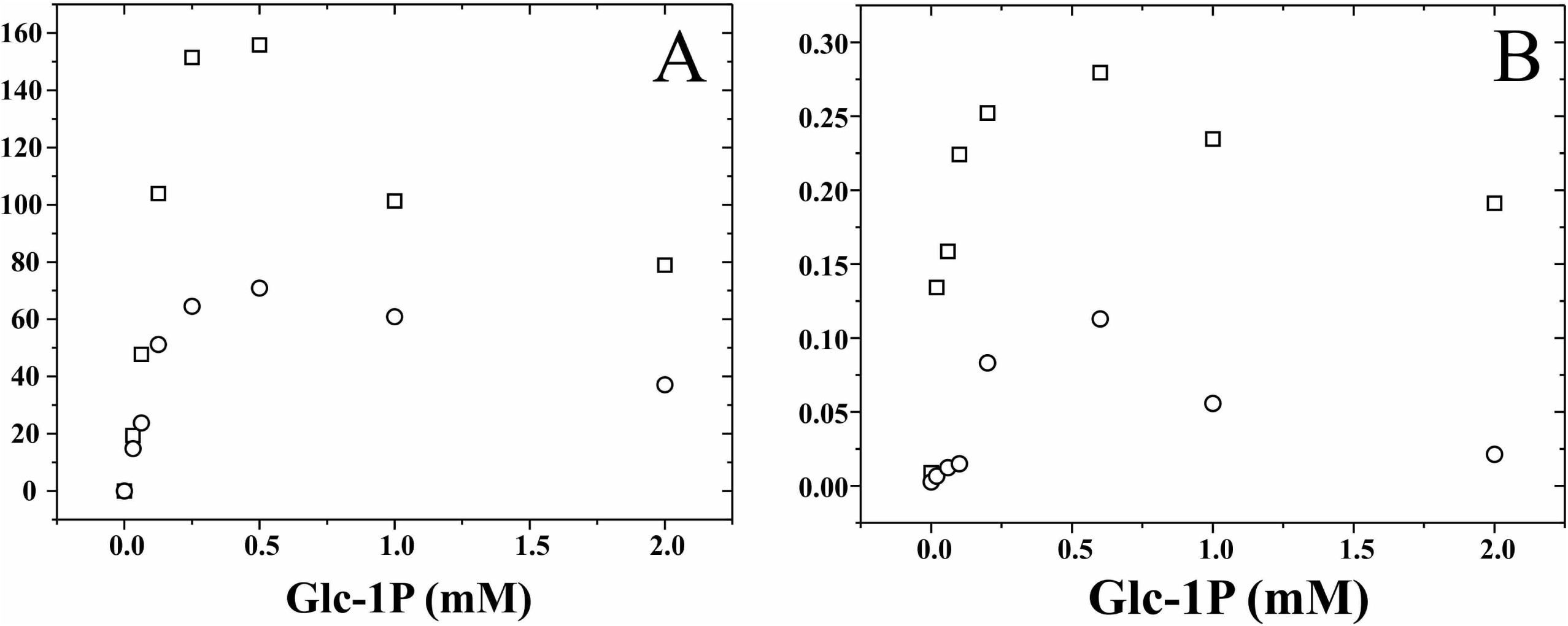
Product inhibition curves in maltose-1P synthesis for *Rjo*GlgAc **(A)** and *Rjo*GlgAb **(B)**. Glc-1P curves were made in presence of 0.3 mM (open circles) or 3 mM (open squares) of ADP-Glc for *Rjo*GlgAc, and with 0.2 mM (open circles) or 1 mM (open squares) for *Rjo*GlgAb.

**Table 2:**
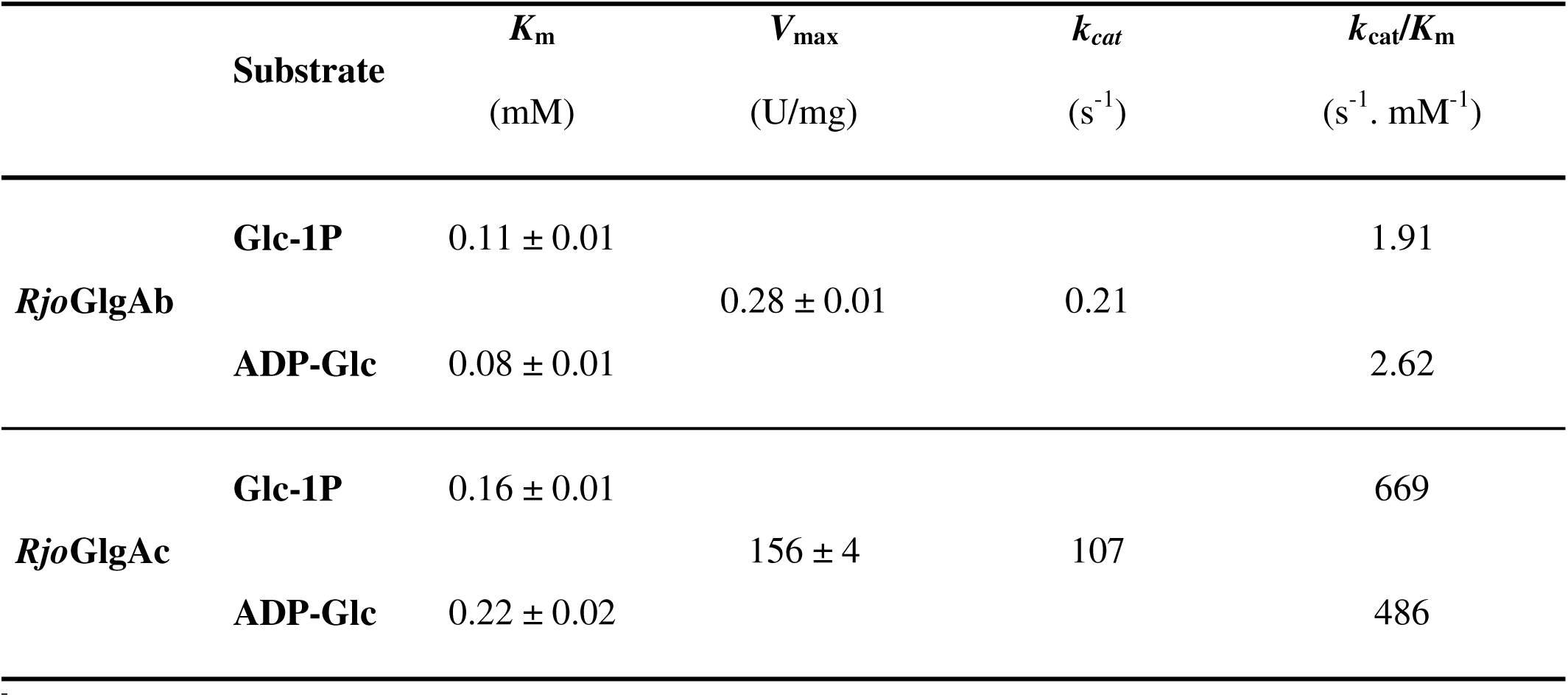
Kinetic parameters of RjoGlgAb and RjoGlgAc for maltose-1P synthesis. k_cat_ were calculated using the corresponding theoretical molecular mass of the monomer.

| Substrate | | $K_{\text{m}}$<br>(mM) | $V_{\text{max}}$<br>(U/mg) | $k_{\text{cat}}$<br>(s<sup>-1</sup>) | $k_{\text{cat}}/K_{\text{m}}$<br>(s<sup>-1</sup> · mM<sup>-1</sup>) |
| --- | --- | --- | --- | --- | --- |
| *RjoGlgAb* | **Glc-1P** | 0.11 ± 0.01 | 0.28 ± 0.01 | 0.21 | 1.91 |
|  | **ADP-Glc** | 0.08 ± 0.01 |  |  | 2.62 |
| *RjoGlgAc* | **Glc-1P** | 0.16 ± 0.01 | 156 ± 4 | 107 | 669 |
|  | **ADP-Glc** | 0.22 ± 0.02 |  |  | 486 |

To determine the *K*_m_ values for *Rjo*GlgAc or *Rjo*GlaAb, we considered the hyperbolic portion of Glc-1P curves. For ADP-Glc plots, the inhibitory effects at high substrate concentrations were absent in the analysis of both enzymes, as shown in Supplementary Figure 5. The *K*_m_ and *k*cat values for *Rjo*GlgAc are in the same order of magnitude as those measured for the *M. tuberculosis* enzyme [31]. This rhodococcal enzyme catalyzes maltose-1P synthesis using the glucosyl-donor ADP-Glc with 124-fold higher catalytic efficiency for Glc-1P than an α-1,4-glucan as acceptors (see Tables 1 and 2). Instead, *Rjo*GlgAb catalyzes both reactions with almost identical *k*_cat_ values, although, notoriously, it showed 13-fold higher efficiency for ADP-Glc utilization in the maltose-1P synthase activity. According to the kinetic parameters shown in Table 1 and Table 2, results suggest that intracellular glycogen synthesis in *R. jostii* may occur via the GlgE pathway, as demonstrated in *M. tuberculosis* [31, 69].

Then, this work constitutes the first report where both actinobacterial GlgA glucosyl-transferases are obtained soluble, actives and comparatively analyzed at the kinetic and structural level. We add novel kinetic information regarding the possibility that another enzyme in the organism, *Rjo*GlgAb, may contribute to the maltosyl donor maltose-1P. Whether this contribution to polyglucan synthesis (glycogen and/or MGLP) has physiological relevance remains to be solved.

### Alternative substrate analysis in maltose-1P synthase activity

Given that both rhodococcal GlgA proteins could catalyze maltose-1P synthesis, we analyzed their ability to use alternative hexoses-1P as aglycon substrates, opening the possibility for a putative synthesis of a heterodisaccaride-1P. Regarding glucosyl-donors, the enzymes were specific for ADP-Glc, and no activity was detected even at a 10 mM UDP-Glc or GDP-Glc and 2 mM Glc-1P (not shown). We also assayed the usage of alternative sugars-1P as glucosyl-acceptors in replacement of Glc-1P. As presented in Figure 4, the *Rjo*GlgAc activity with most sugar-1P analyzed (at 2 mM) is about 10% *versus* the one measured with Glc-1P. However, the enzyme was highly active when GlcN-1P was tested, reaching a relative activity of 40% of that obtained with 2 mM Glc-1P. *Rjo*GlgAb also showed its highest activity when Glc-1P was the acceptor, and all alternative sugar-1Ps assayed showed activities lower than 20% when compared to Glc-1P (Figure 4). Given the *Rjo*GlgAb scarce activity values for maltose-1P synthesis and that the use of other sugar-1P were even lower, our kinetic characterization was not further since the physiological relevance of these alternative activities would be null. However, it should be considered that the potential of both rhodococcal GlgAs enzymes to produce heterodisaccarides-1P constitutes the basis for developing new biocatalytic tools usable for cell-free glycobiology strategies.

**Figure 4:**
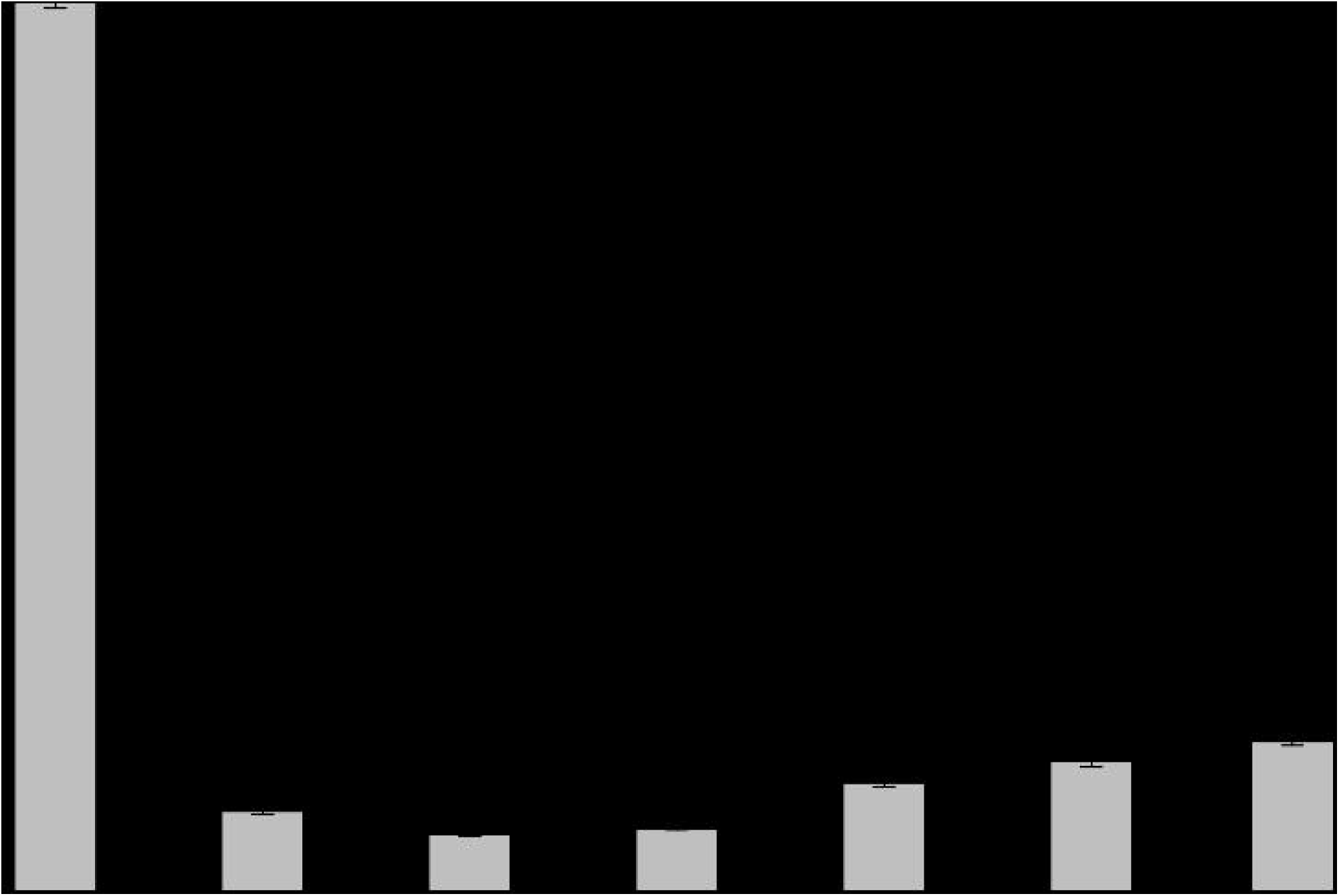
Activity of *Rjo*GlgAb **(grey)** and *Rjo*GlgAc **(black)** using alternative sugars-1P. Histogram shows the relative activities obtained with different sugars-1P assayed at 2 mM and 1 mM ADP-Glc. The value of 1 belongs to activities of 0.28 U/mg (*Rjo*GlgAb) and 156 U/mg (*Rjo*GlgAc) when using Glc-1P as a substrate.

We proceeded with the *Rjo*GlgAc kinetic characterization regarding GlcN-1P utilization (Figure 5). The amino sugar-1P substrate showed a *k*_cat_ of 36.7 s^-1^ and a *K*_m_ value of 2.2 ± 0.1 mM for this substrate (at 1 mM ADP-Glc). Strikingly, the GlcN-1P curves did not exhibit the inhibitory effect at higher substrate concentrations, depicting a behavior that deviates slightly from hyperbolic (n_H_ =1.2). After analyzing ADP-Glc curves obtained at 1 mM GlcN-1P, we calculated a *K*_m_ value of 0.41 ± 0.06 mM. The catalytic efficiency (as of *k*_cat_/*K*_m_) performed by *Rjo*GlgAc with GlcN-1P, although 40-fold lower than for Glc-1P (see Table 2), is higher than the corresponding efficiency as a glucan elongating enzyme. Still, kinetic values suggest that GlcN-1P might be metabolically relevant. In this context, the high efficiency of *Rjo*GlgAc towards GlcN-1P reinforces the putative importance of this metabolite, being directed to yet unknown intracellular fates, different for being just a mere intermediary between central metabolism and peptidoglycan synthesis/degradation [20,29,30]. Worthy to note the *Rjo*GlgAc apparent affinity for GlcN-1P is in the same millimolar order as that of the rhodococcal GlgCs for the same substrate, as reported earlier [29]. In addition, we extended the study to other bacterial ADP-GlcPPases and determined that GlcN-1P acts as a secondary yet efficient substrate in most cases, concomitantly with activation by GlcN-6P [30]. Taken together, we are tempted to speculate that in certain organisms (such as *R. jostii*) and conditions, GlcN might find a way to be incorporated into different molecules.

**Figure 5:**
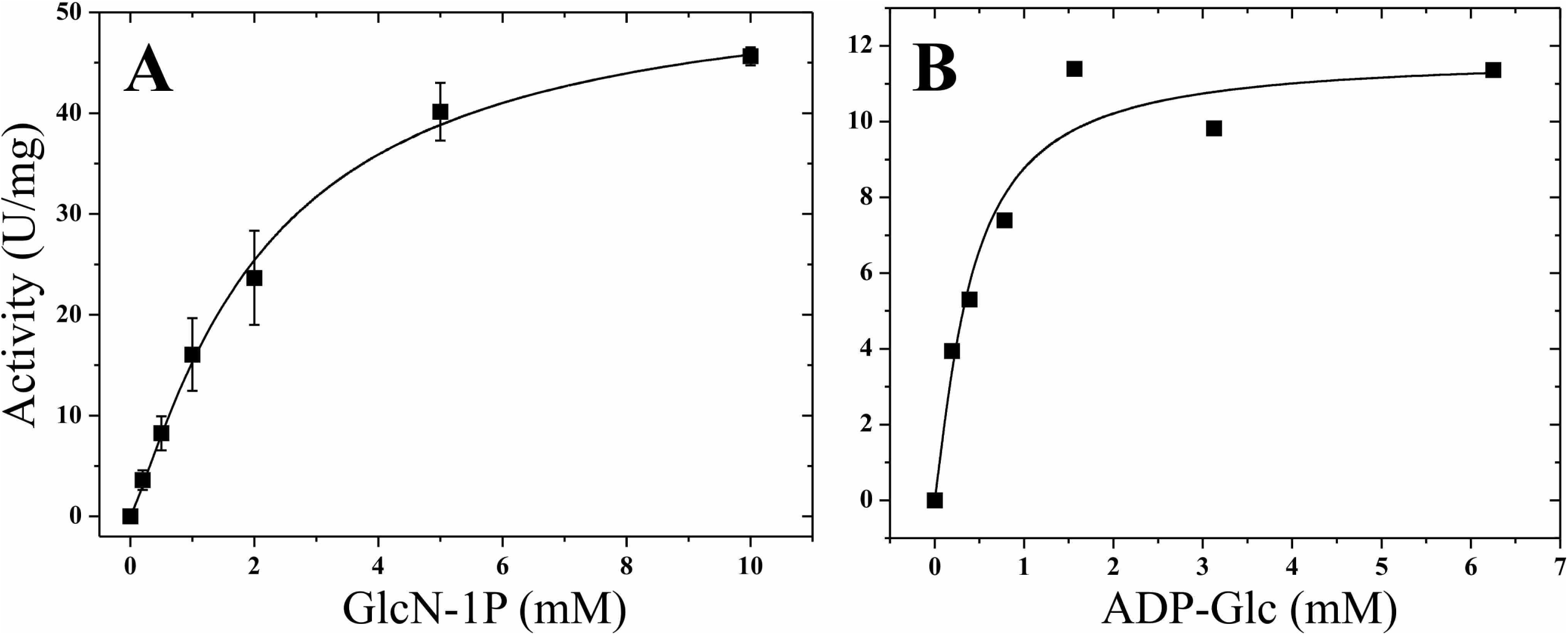
Saturation curves for maltose-1P synthesis of *Rjo*GlgAc with GlcN-1P (A) and ADP-Glc (B). 1 mM of ADP-Glc was used for GlcN-1P saturation curve, as well 1 mM GlcN-1P for ADP-Glc saturation curves.

## Discussion

Glycogen and starch synthases were grouped into the glycosyltransferase families GT3 and GT5 according to the Carbohydrate Active Enzyme database (CAZy) [71]. The former includes glycogen synthases, mainly from fungi and mammals using UDP-Glc as the preferable glucosyl donor (EC 2.4.1.11) and being allosterically regulated [72–74]. Instead, the GT5 family comprises glycogen and starch synthases from bacteria and plants, respectively, possessing ADP-Glc as the specific glucosyl-donor (EC 2.4.1.21), with no reports regarding allosteric regulation so far [75–78]. In bacteria and plants, the key step of glucan elongation is at the level of ADP-Glc synthesis by the allosteric, highly regulated ADP-GlcPPase (EC 2.7.7.27) [28, 79]. On the other hand, GT4 glycosyltransferases compose one of the largest families among CAZy classification, possessing enzymes with diverse functions, catalyzing a wide repertory of reactions and, consequently, displaying differences in their specificity towards glycosyl donors and/or acceptors [54, 71]. To date, the family accounts for 252,202 members, with 160 characterized enzymes and 35 solved three-dimensional structures [71].

Recently, a new activity, ADP-Glc dependent α-maltose-1-phosphate synthase (EC 2.4.1.342) was incorporated into the GT4 family after the report of the mycobacterial GlgM enzyme [31], which was crystalized [32]. Previously, mycobacterial GlgM was referred to as GlgA as of the typical bacterial GT5 glycogen synthases (EC 2.4.1.21), after its relatively high identity with those enzymes (about 40%). The mycobacterial GlgA was already characterized in its “classical” GlgA activity [41], similar to the GlgA enzyme from *S. coelicolor* [57]. In addition, in *M. tuberculosis* and *M. smegmatis* the gene encoding GlgM locates adjacent to *glgC*, which codifies the ADP-GlcPPase from the classical glycogen biosynthesis pathway [80, 81]. Likewise, other actinobacterial members such as *R. jostii*, *R. fascians*, and *S. coelicolor*, show the same synteny for the *glgC*/*glgA* (*glgM*) genes location [5,82,83]. Indeed, to differentiate each GlgA studied in this work, we named GlgAc to the glycosyl-transferase whose encoding gene (*glgA*) is next to the one coding for ADP-GlcPPase (*glgC*) in *R. jostii*. We called GlgAb the other glucosyl-transferase because its gene was immediately neighboring the one encoding a putative glucan branching enzyme (usually referred to as GlgB) [5]. *Rjo*GlgAb is highly homolog to Rv3032, a putative glucosyl-α-1,4-transferase described to participate in the elongation of MGLPs molecules in members of the *Mycobacterium* genus [26,56,63,67]. However, besides several biochemical reports, the enzyme was never kinetically characterized and its activity and/or substrate preference were inferred from results with mycobacterial mutants [56, 63].

Until the work presented here, the only GlgM kinetic characterizations were from mycobacterial sources [31, 32]. Also, the only accounts for Rv3032 function were from *M. tuberculosis* [63, 67]. The basis behind the understanding of these mycobacterial glycosyl-transferases was (mostly) the search for anti-tuberculosis targets [26, 84]. Certainly, the maltosyl-transferase GlgE (EC 2.4.99.16) related to polyglucan synthesis, which uses as substrate the maltose-1P produced by GlgM, was postulated as an anti-tuberculosis drug target. GlgE inhibition leads to a self-poisoning accumulation of maltose-1P [68, 84]. The GlgE biochemical role was later approached in other organisms, such as *Chlamydia* [85] and *Pseudomonas* [86]. Nevertheless, the GlgE involvement on carbon partitioning in biotechnological relevant species (*e*.*g*., biofuel and/or value-added biomolecule production) remains uncovered. In agreement with this scenario, it still lacks GlgM characterizations different from a mycobacterial source. Then, advancing in describing new GlgM enzymes is imperative, not only to better understand glycogen metabolism in Actinobacteria but to reveal hints regarding the evolutionary history of the maltose-1P synthase activity. In this regard, we approached the study of glycogen synthesis-related enzymes in the oleaginous bacterium *R. jostii*, which sums up our previous reports regarding rhodococcal ADP-GlcPPases regulatory properties [24, 29]. In *R. jostii*, glycogen behaves as a temporal carbon storage molecule connected to lipid synthesis. *Ergo*, further knowledge concerning the specific properties of glycogen biosynthetic enzymes in *Rhodococci* is crucial to harness or improve their properties as biofactories producing biofuel precursors, such as TAGs [10].

The work presented here provides kinetic data to understand polysaccharide synthesis in Actinobacteria, particularly intracellular glycogen production in *R. jostii*. We demonstrate that *Rjo*GlgAc, predicted to be a “classical GlgA”, is a maltose-1P synthase (EC 2.4.1.342) with two orders of magnitude higher *k*_cat_ than the glycogen synthase activity (EC 2.4.1.21). The enzyme uses ADP-Glc as the specific glucosyl donor to glycosylate Glc-1P more efficiently than glycogen. In agreement with recent reports regarding mycobacterial GlgA protein [31, 32], *Rjo*GlgAc should be ascribed as GlgM in future works. Worthy to remark, we report here that *Rjo*GlgAb also possesses maltose-1P activity, although in this enzyme, Glc-1P is a glycosyl-acceptor 350-fold less efficient than in *Rjo*GlgAc. Then, results strongly suggest that carbon for glycogen synthesis in *R. jostii* would flow through maltose-1P *via* the GlgE pathway [69, 70], being *Rjo*GlgAc the responsible enzyme in synthesizing the substrate for glucan elongation by GlgE.

The maltosyl-transferase GlgE was first described in *M. tuberculosis* by Alan D. Elbein, unfolding the last enzymatic step in the pathway converting trehalose into glycogen [69]. Later, bioinformatic studies demonstrated that the GlgE pathway was present in up to 14% of the bacterial genomes available at that time [70]. The *R. jostii* genomic information evidences a *glgE* gene that encodes the respective maltosyl-transferase GlgE. The latter shows a 65% identity regarding the enzyme from *M. tuberculosis* [68, 69] and possesses the critical amino acidic residues for activity postulated after the crystallization and solved structure of the GlgE from *S. coelicolor* [87, 88] (not shown). Taken together, *Rjo*GlgAc kinetic results sustain the idea of a functional GlgM-GlgE pathway in *R. jostii* where intracellular glycogen would be elongated in two glucosidic moieties [69]. In accordance, it was suggested that those organisms possessing both classical and GlgE glycogen synthesis pathways use the latter to produce the glucan [66]. In this context, this work adds new elements (*Rjo*GlgAc) to be considered in the interplay between glycogen and lipid metabolisms in *R. jostii*. Besides, the classical GlgA activity depicted by *Rjo*GlgAc, where the glucan is the acceptor of one glucose unit from ADP-Glc, showed kinetic efficiency values which are in the range of metabolic feasibility [64, 65]. Remarkably, the apparent affinity for glycogen in the *Rjo*GlgAc is among the highest reported; whether the relevance of this classical pathway in the *in vivo* glycogen accumulation remains to be approached.

The kinetic and comparative characterization presented herein provides a new enzymatic actor, *Rjo*GlgAb, to the discussion regarding maltose-1P synthesis, in addition to the *Rjo*GlgAc described above and maltokinase (EC 2.7.1.175). Maltokinases phosphorylate maltose employing NTP (mainly ATP) as the phosphoryl donor, as described in studies with the enzyme from actinobacterial sources [89–93]. Genomic *R. jostii* analysis allowed us to identify one gene encoding a putative maltokinase [5]. The *Rjo*GlgAb maltose-1P synthase activity is significantly lower than in *Rjo*GlgAc, but still, its kinetic efficiency values suggest a metabolic probability [64]. The physiological relevance should be addressed to confirm each enzyme’s contribution to the intracellular pool of maltose-1P.

We show that *Rjo*GlgAb depicts both activities, glycogen and maltose-1P synthases, with almost identical *k*_cat_ values (presented in Table 1 and Table 2). Since results showed that the rhodococcal GlgM is *Rjo*GlgAc (see above), we focused on analyzing the classical glycogen synthase activity in *Rjo*GlgAb. We used glycogen from rabbit muscle, the commonly available substrate used to characterize several glycogen and starch synthases [41,48,49,57,76,94]. Kinetic results indicate that *Rjo*GlgAb catalyzes the transfer of a glycosidic moiety to glycogen, preferring ADP-Glc as the sugar donor. The activity was in the same order of magnitude that actinobacterial analogs characterized before [41, 57], although lower than in “model” glycogen synthases, such as those from *E. coli* [48, 50] and *A. tumefaciens* [47]. We observed that glycogen from oyster produced higher *Rjo*GlgAb activities, indicating a preference for short α-1,4-glucan branches, which could be as low as 2-3 units in this type of glucan [58, 59]. Shorter branches are also present in immature MGLP molecules, postulated as possible substrates for mycobacterial Rv3032 [26, 67]. Thus, MGLP could be hypothesized as a putative substrate for *Rjo*GlgAb. When glycogen from (actinobacterial) sources phylogenetically related to *R. jostii* was analyzed, *Rjo*GlgAb displayed higher glucan affinities (around 10^-2^ mg/ml) in the same order of magnitude than *Rjo*GlgAc affinity towards glycogen.

Then, it can be assumed that the GlgA GT4 type protein evolved conserving the ability to efficiently bind to glucan molecules, despite being structurally different from GT5 bacterial glycogen synthases [51]. In the particular case of *Rjo*GlgAb, the low “glycogen synthase” activity enzyme could be ascribed to being analyzed as an alternative substrate or omitted some activating factor yet to be identified. At this point, we hypothesized that a glucosyl-glycerate derivative and/or an immature MGLP molecule would act as an aglycon for the glucosyl-transferase activity, as proposed for the mycobacterial homolog Rv3032 [26,63,67]. In this regard, we analyzed the *R. jostii* genomic information looking for genes encoding enzymes associated with glucosyl-glycerate metabolism and MGLPs biosynthesis.

We found that *R. jostii* encodes putative GpgS (EC 2.4.1.266) and GpgP (EC 3.1.3.85) enzymes, which respectively catalyze the synthesis and hydrolysis of glucosyl-3-phosphoglycerate to produce glucosyl-glycerate, the precursor of MGLP synthesis [26]. The presence of a *ggH* gene in *Rhodococcus* was also reported, which probably plays a regulatory role in MGLP synthesis [26,95,96]. Also, *R. jostii* would produce an OctT protein[97], which catalyzes the production of octanoyl-diacylglycerate, which may help recruit the subsequent enzymes in the MGLP synthetic pathway, such as Rv3032 that elongates the glucan [56, 67]. This preliminary analysis supports the idea of a functional metabolism for glucosyl-glycerate and MGLPs in the organism. MGLP are known to regulate fatty acid metabolism *in vitro* [26, 27], then opening a new opportunity to explore the link between carbohydrate and lipid metabolisms and a probable impact on TAGs production for biodiesel purposes [10].

Except for a cyanobacterium case [53], almost no reports regarding duplicated glycogen synthases are available in the bibliography. The duplicated GlgA-like proteins were noticed in Mycobacteria, but the kinetic analysis was analyzed only for one of those which resulted in the description of a new type of activity, maltose-1P synthase [31]. To understand the basis for this “duplication” we constructed a phylogenetic tree with GT4 GlgA sequences sharing an identity to *Rjo*GlgAb higher than 40% (the identity between *Rjo*GlgAb and *Rjo*GlgAc). For the sake of comparative analysis, we added representative GlgMs, GT5 bacterial glycogen synthases (GlgA) and GT3 starch synthases, as presented in the phylogenetic tree shown in Figure 6. The major output from the analysis relies on the fact that we found *Rjo*GlgAb hits limited to the actinobacterial groups *Streptomyces*, *Micrococcus*, *Bifidobacterium*, *Corynebacterium*/*Gordonia*, *Rhodococcus* and *Mycobacterium*. No other bacterial representatives were found. Remarkably, *Rhodococcus* and *Mycobacterium* constitute the closest branches, reinforcing the idea that kinetic results presented here for *Rjo*GlgAb could help in inferring regarding the enzymatic behavior of the mycobacterial Rv3032.

**Figure 6:**
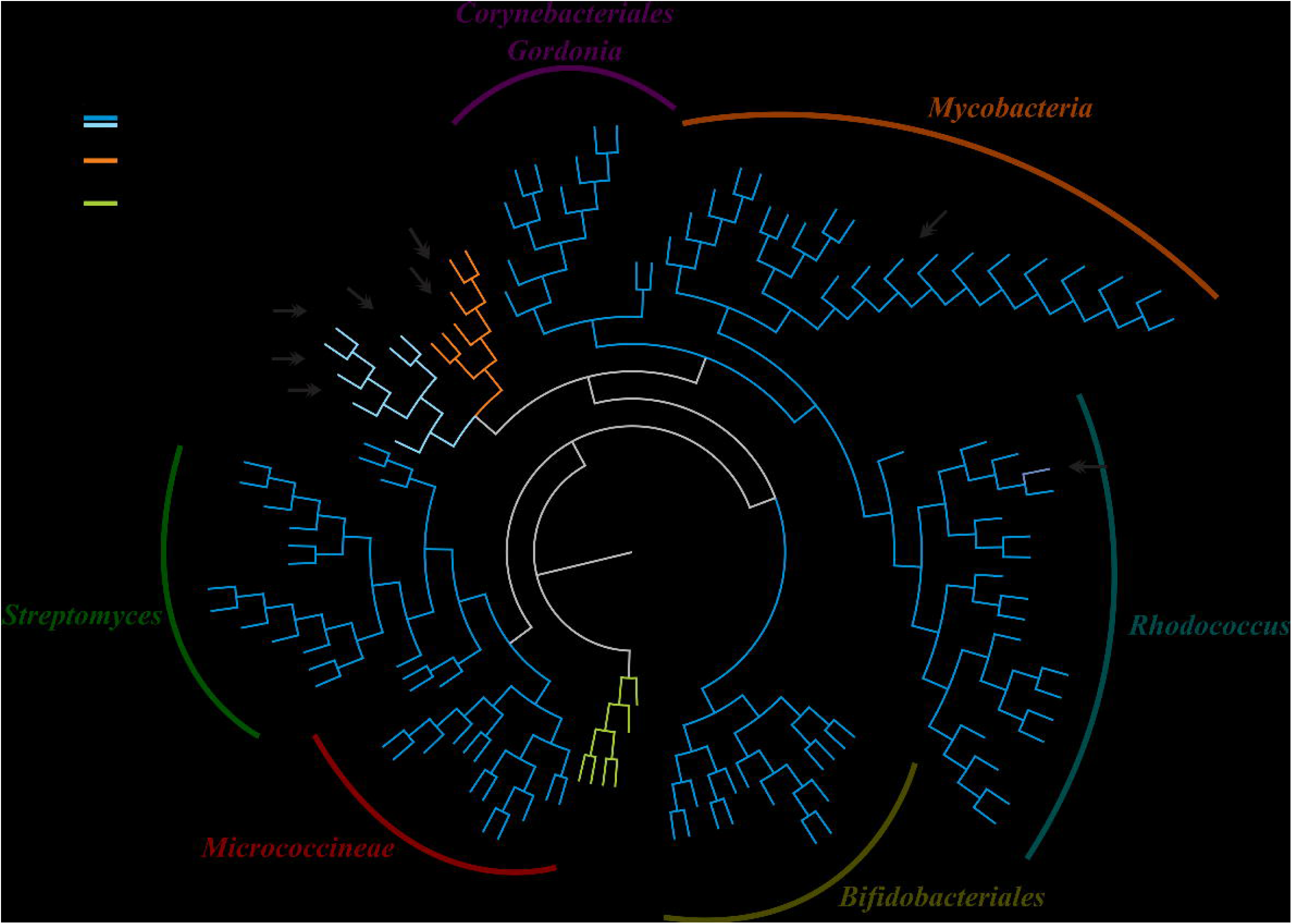
Phylogenetic analysis from different glycogen synthases. Sequences from actinobacterial GlgAs belonging to GT3, GT4 and GT5 glycosyl-transferases groups were collected and the tree was constructed as described at Materials and Methods section. The numerical code assigned to each sequence is indexed at Supplemental table 2. The structurally and/or kinetically characterized enzymes that are taken as reference are marked, and different bacterial genera are delimited.

Next to *Rhodococcus* and *Mycobacterium* locates the *Corynebacterium* group, which has an average identity of 62% and 64% with the mentioned proteins (*Rjo*GlgAb and Rv3032, respectively). Figure 6 shows that members of *Bifidobacterium* situate farther, in an almost independent branch. The average identity with *Rjo*GlgAb and Rv3032 of the *Bifidobacterium* proteins is 20% and 27%, respectively. Yet, this *Bifidobacterium* clade locates separated from each other group of glycosyl-transferase presented in the phylogenetic tree. Also, the target proteins from *Streptomyces* and *Micrococcus* showed identity values between 31% and 32% compared to *Rjo*GlgAb and placed in the phylogenetic tree more distantly (Figure 6). On the other hand, GlgM enzymes grouped with the GT5 GlgAs while the GT3 enzymes formed an independent clade. In a complete view, the *Rjo*GlgAb/Rv3032 type of enzymes seems to be present in actinobacterial members, coincident with MGLP description or isolation, mostly limited to species from *Nocardia*, *Streptomyces* and *Mycobacterium* [26,56,67].

Our results prompt us to focus on other actinobacterial members with biotechnological relevance and continue elucidating the occurrence, metabolism and possible role of MGLP. As well, given the high identity between the *Rjo*GlgAb-like proteins shown in Figure 6, the particular structural features (super-oligomeric state) described for *Rjo*GlgAb (absent in other GT4, GT5 and GT3 GlgA type enzymes), the common glucan elongation activity shared for the three GT families, and the maltose-1P activity occurring in both type of actinobacterial GT4 GlgAs, a scenario for elucidate the evolutionary aspects behind these properties has been set in this work. The kinetic and structural characterization of other *Rjo*GlgAb homologs from the above-mentioned actinobacterial groups will be vital to achieving this knowledge.

Another key result from the study of the rhodococcal GlgAs is the ability shown by *Rjo*GlgAc in using GlcN-1P as an aglycon. Our group previously proposed that the hexosamine-1P may have an alternative metabolic fate, different from a mere intermediary between primary metabolism and peptidoglycan synthesis [20, 29]. Indeed, the catalytic efficiency showed by *Rjo*GlgAc for GlcN-1P is in the same order of magnitude as the specific for GlcN-1P pyrophosphorylase *Rjo*GalU2 (∼59 mM^-1^ seg ^-1^) [20] and is higher than those obtained for rhodococcal ADP-GlcPPases, which are around 0.6 mM^-1^ seg ^-1^, after activation by Glc-6P and/or GlcN-6P [29, 30]. These efficiency parameters suggest a metabolic plausibility [64]. Thus, the enzymatic information presented here reinforces the hypothesis that rhodococcal glycogen-related enzymes (ADP-GlcPPase and *Rjo*GlgAc) can develop a secondary activity with GlcN-1P. Then *Rjo*GlgAc could be considered as a new biocatalyst in the GlcN-1P node, supporting the idea of a partitioning node at the level of the hexosamine-1P, similar to that for Glc-1P. Our current focus relies on determining if the ability of rhodococcal enzymes to channel glucosamine moieties *in vivo* to different molecules is metabolically functional or if it remains as part of the underground metabolism [98].

The work presented herein with two bifunctional GlgA-like enzymes sharing the same activities, although to a different extent and with remarkable structural differences, may constitute a molecular hint for future studies related to their structure-to-function relationships. Our kinetic characterizations together with bioinformatic analysis [70] and biochemical assays already available with mycobacterial GlgA (GlgM) and Rv3032 mutants[63], are a cornerstone to unwrap evolutionary aspects of GT4 “glycogen synthases’’ appearance. Also, to understand how this “specialization” is related to bacterial GT5 glycogen synthases, and/or the co-existence with the metabolism of carbohydrate molecules such as glycogen, MGLP, maltose, trehalose and glucosyl-glycerate.

## Supporting information

Supplementary material Rhodococcal GlgAs

## Acknowledgements

AAI, HMA and MDAD are career investigator members of the Consejo Nacional de Investigaciones Científicas y Técnicas (CONICET), Argentina. AEC and MVF are fellowship holders from CONICET. This work was supported by grants from ANPCyT (PICT-2018-00929 and PICT-2020-03326 to AAI; and PICT’18 00698 to MDAD) and CONICET (PIP2015-2016 0529 to HMA).

