## Supplementary material Rhodococcal GlgAs for "Comparative analysis between two GT4 glycosyltransferases related to polysaccharide biosynthesis in *Rhodococcus jostii* RHA1"

**<u>Supplemental Table 1</u>:** Primers used for genes amplification. In bold are the restriction enzyme cut site then used for cloning.

| **Primer** | **Sequence** | **Site** | **Tm (°C)** |
|---|---|---|---|
| *RjoglgAb*fow | 5'-**CATATG**AAGATCCTGATCGTGTCGTGGGAATAC-3' | *Nde*I | 63.0 |
| *RjoglgAb*rev | 5'-**GAGCTC**TCAGCGACCGGGCAGCG-3' | *Sac*I | 72.1 |
| *RjoglgAc*fow | 5'-**CATATG**CGGGTGGCGATGATGACCAAGG-3' | *Nde*I | 67.4 |
| *RjoglgAc*rev | 5'-**AAGCTT**TCAGCGCCGGGACAGTGCG-3' | *Hind*III | 70.7 |

**<u>Supplemental Table 2</u>:** Number codes for sequences used to build the phylogenetic tree (showed in Figure 6).

| | |
|---|---|
| **1** | Rhodococcus jostii GlgAb |
| **2** | WP_005259541.1 glycosyltransferase family 4 protein [Rhodococcus opacus] |
| **3** | WP_009479652.1 glycosyltransferase family 4 protein [Rhodococcus sp. JVH1] |
| **4** | WP_037245393.1 glycosyltransferase family 4 protein [Rhodococcus wratislaviensis] |
| **5** | WP_042951480.1 glycosyltransferase family 4 protein [Rhodococcus erythropolis] |
| **6** | WP_092800949.1 glycosyltransferase family 4 protein [Rhodococcus globerulus] |
| **7** | WP_210378328.1 glycosyltransferase family 4 protein [Rhodococcus qingshengii] |
| **8** | WP_064077610.1 glycosyltransferase family 4 protein [Rhodococcus hoagii] |
| **9** | WP_127914358.1 glycosyltransferase family 4 protein [Rhodococcus agglutinans] |
| **10** | WP_031938991.1 glycosyltransferase family 4 protein [Rhodococcus defluvii] |
| **11** | WP_142097067.1 glycosyltransferase family 4 protein [Rhodococcus spelaei] |
| **12** | WP_072750217.1 glycosyltransferase family 4 protein [Rhodococcus maanshanensis] |
| **13** | WP_136906490.1 glycosyltransferase family 4 protein [Rhodococcus oryzae] |
| **14** | WP_127950893.1 glycosyltransferase family 4 protein [Rhodococcus xishaensis] |
| **15** | WP_068156674.1 glycosyltransferase family 4 protein [Rhodococcus phenolicus] |
| **16** | WP_060651425.1 glycosyltransferase family 4 protein [Rhodococcus pyridinivorans] |
| **17** | WP_138997446.1 glycosyltransferase family 4 protein [Rhodococcus zopfii] |
| **18** | WP_094980283.1 glycosyltransferase family 4 protein [Rhodococcus pyridinivorans] |
| **19** | WP_032403133.1 glycosyltransferase family 4 protein [Rhodococcus fascians] |
| **20** | WP_119699867.1 glycosyltransferase family 4 protein [Rhodococcus ruber] |
| **21** | WP_155377522.1 glycosyltransferase [Cellulomonas sp. JZ18] |
| **22** | WP_136520107.1 glycosyltransferase [Cellulomonas telluris] |
| **23** | WP_143418856.1 glycosyltransferase [Georgenia sp. Z446] |
| **24** | WP_221655739.1 glycosyltransferase [Actinotalea ferrariae] |
| **25** | WP_159798431.1 glycosyltransferase [Puerhibacterium puerhi] |
| **26** | WP_175009829.1 glycosyltransferase [Cellulosimicrobium sp. TH-20] |
| **27** | WP_111251743.1 glycosyltransferase [Xylanimonas oleitrophica] |
| **28** | WP_208195003.1 glycosyltransferase [Cellulomonas sp. zg-ZUI157] |
| **29** | WP_069984692.1 glycosyltransferase [Isoptericola variabilis] |
| **30** | WP_168946323.1 glycosyltransferase [Cellulosimicrobium aquatile] |

| | |
|---|---|
| **31** | WP_218870505.1 glycosyltransferase [Herbiconiux flava] |
| **32** | WP_171108803.1 glycosyltransferase [Promicromonospora citrea] |
| **33** | WP_034228905.1 glycosyltransferase, partial [Actinotalea ferrariae] |
| **34** | WP_152230542.1 glycosyltransferase [Georgenia ruanii] |
| **35** | WP_123045853.1 glycosyltransferase [Cryobacterium tepidiphilum] |
| **36** | WP_157427622.1 glycosyltransferase [Agromyces salentinus] |
| **37** | WP_018385493.1 glycosyltransferase [Streptomyces vitaminophilus] |
| **38** | WP_093600331.1 glycosyltransferase [Streptomyces jietaisiensis] |
| **39** | WP_185393590.1 glycosyltransferase [Streptomyces rutgersensis] |
| **40** | WP_052863203.1 glycosyltransferase [Streptomyces niger] |
| **41** | WP_190131612.1 glycosyltransferase [Streptomyces mashuensis] |
| **42** | WP_093656013.1 glycosyltransferase [Streptomyces radiopugnans] |
| **43** | WP_006143302.1 glycosyltransferase [Streptomyces griseoaurantiacus] |
| **44** | WP_101257480.1 glycosyltransferase [Streptomyces barkulensis] |
| **45** | WP_086705093.1 glycosyltransferase [Streptomyces variabilis] |
| **46** | WP_155072438.1 glycosyltransferase [Streptomyces taklimakanensis] |
| **47** | WP_189943392.1 glycosyltransferase [Streptomyces aurantiogriseus] |
| **48** | WP_062046197.1 glycosyltransferase [Streptomyces canus] |
| **49** | WP_146469256.1 glycosyltransferase [Streptomyces misionensis] |
| **50** | WP_093849588.1 glycosyltransferase [Streptomyces pini] |
| **51** | WP_200305176.1 glycosyltransferase [Streptomyces adelaidensis] |
| **52** | WP_093841954.1 glycosyltransferase [Streptomyces harbinensis] |
| **53** | WP_151167338.1 glycosyltransferase [Streptomyces albidoflavus] |
| **54** | WP_085180693.1 glycosyltransferase family 4 protein [Mycobacterium bohemicum] |
| **55** | WP_005102402.1 glycosyltransferase family 4 protein [Mycobacteroides abscessus] |
| **56** | WP_134070606.1 glycosyltransferase family 4 protein [Mycobacteroides salmoniphilum] |
| **57** | WP_064629781.1 glycosyltransferase family 4 protein [Mycobacteroides immunogenum] |
| **58** | WP_070909840.1 glycosyltransferase family 4 protein [Mycobacteroides saopaulense] |
| **59** | WP_064927525.1 glycosyltransferase family 4 protein [Mycolicibacterium elephantis] |
| **60** | WP_067827337.1 glycosyltransferase family 4 protein [Mycobacterium mantenii] |
| **61** | WP_087078554.1 glycosyltransferase family 4 protein [Mycobacterium dioxanotrophicus] |

| | |
|---|---|
| **62** | WP_055112676.1 glycosyltransferase family 4 protein [Mycolicibacterium peregrinum] |
| **63** | WP_075234739.1 glycosyltransferase family 4 protein [Mycobacterium colombiense] |
| **64** | WP_200995943.1 glycosyltransferase family 4 protein [Mycobacteroides chelonae] |
| **65** | WP_110917301.1 glycosyltransferase family 4 protein [Mycobacterium holsaticum] |
| **66** | WP_083162078.1 glycosyltransferase family 4 protein [Mycobacterium aquaticum] |
| **67** | WP_083066435.1 glycosyltransferase family 4 protein [Mycobacterium arosiense] |
| **68** | WP_069395022.1 glycosyltransferase family 4 protein [Mycobacterium shimoidei] |
| **69** | WP_077098994.1 glycosyltransferase family 4 protein [Mycobacterium terramassiliense] |
| **70** | WP_064936680.1 glycosyltransferase family 4 protein [Mycobacterium intracellulare] |
| **71** | WP_085077362.1 glycosyltransferase family 4 protein [Mycobacterium palustre] |
| **72** | WP_038435961.1 glycogen synthase [Mycobacterium tuberculosis] Mtb3032 |
| **73** | WP_117414877.1 glycosyltransferase family 4 protein [Mycobacterium marinum] |
| **74** | WP_023865110.1 glycosyltransferase family 4 protein [Mycobacterium avium] |
| **75** | WP_068970700.1 glycosyltransferase family 4 protein [Corynebacteriales] |
| **76** | WP_040518439.1 glycosyltransferase family 4 protein [Gordonia aichiensis] |
| **77** | WP_127962726.1 glycosyltransferase family 4 protein [Gordonia alkanivorans] |
| **78** | WP_040515947.1 glycosyltransferase family 4 protein [Gordonia amarae] |
| **79** | WP_170193889.1 glycosyltransferase family 4 protein [Gordonia asplenii] |
| **80** | WP_012834984.1 glycosyltransferase family 4 protein [Gordonia bronchialis] |
| **81** | WP_161927790.1 glycosyltransferase family 4 protein [Gordonia crocea] |
| **82** | WP_059039212.1 glycosyltransferase family 4 protein [Gordonia desulfuricans] |
| **83** | WP_164309046.1 glycosyltransferase family 4 protein [Gordonia hankookensis] |
| **84** | WP_124710955.1 glycosyltransferase family 4 protein [Gordonia insulae] |
| **85** | WP_040534022.1 glycosyltransferase family 4 protein [Gordonia rhizosphera] |
| **86** | WP_158216449.1 glycosyltransferase [Bifidobacterium hapali] |
| **87** | WP_052119042.1 glycosyltransferase [Bifidobacterium callitrichos] |
| **88** | WP_206666338.1 glycosyltransferase [Bifidobacterium jacchi] |
| **89** | WP_225431778.1 glycosyltransferase [Bifidobacterium platyrrhinorum] |
| **90** | WP_129859801.1 glycosyltransferase family 4 protein [Bifidobacterium pseudolongum] |
| **91** | WP_123644384.1 glycosyltransferase family 4 protein [Bifidobacterium mongoliense] |
| **92** | WP_101428719.1 glycosyltransferase family 4 protein [Bifidobacterium pseudolongum] |

| | |
|---|---|
| **93** | WP_137656081.1 glycosyltransferase [Bifidobacterium moukalabense] |
| **94** | WP_168984540.1 glycosyltransferase [Bifidobacterium thermophilum] |
| **95** | WP_051867145.1 glycosyltransferase [Bifidobacterium crudilactis] |
| **96** | WP_094664272.1 glycosyltransferase [Bifidobacterium tissieri] |
| **97** | WP_209164051.1 glycosyltransferase [Bifidobacterium dentium] |
| **98** | WP_152354779.1 glycosyltransferase family 4 protein [Bifidobacterium apri] |
| **99** | WP_125968241.1 glycosyltransferase family 4 protein [Bifidobacterium samirii] |
| **100** | WP_045935485.1 glycosyltransferase family 4 protein [Bifidobacterium asteroides] |
| **101** | Rhodococcus jostii GlgAc |
| **102** | Kocuria rhizophila GlgA |
| **103** | S. coelicolor GlgA |
| **104** | M. tuberculosis GlgA |
| **105** | Corynebacterium glutamicum GlgA |
| **106** | M. smegmatis GlgM |
| **107** | M. tuberculosis GlgM |
| **108** | E. coli GlgA |
| **109** | Nitrosomonas europaea GlgA |
| **110** | A. tumefaciens GlgA |
| **111** | Arabidopsis thaliana GlgA |
| **112** | Prunus persica GlgA |
| **113** | Pseudomonas aeruginosa GlgA |
| **114** | Prevotella intermedia GlgA |
| **115** | Saccharomyces cerevisiae GlgA1 |
| **116** | Saccharomyces cerevisiae GlgA2 |
| **117** | Giardia lamblia GlgA |
| **118** | Homo sapiens GlgA1 |
| **119** | Homo sapiens GlgA2 |

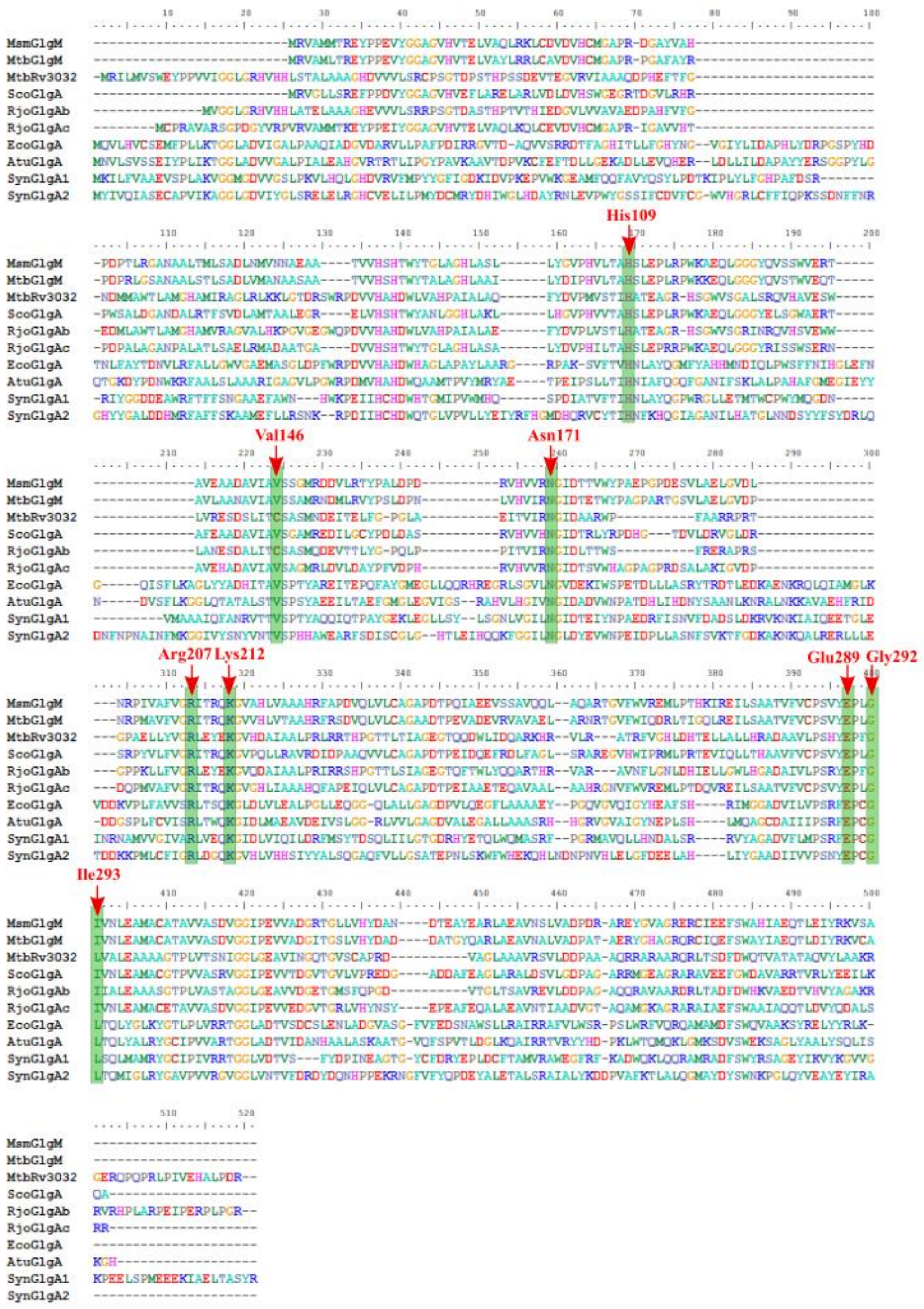


**Supplemental Figure 1:** Sequence alignment of glycogen synthases from different microorganisms. The highlighted residues correspond to key binding and catalytic amino acids in GlgA and GlgM proteins.

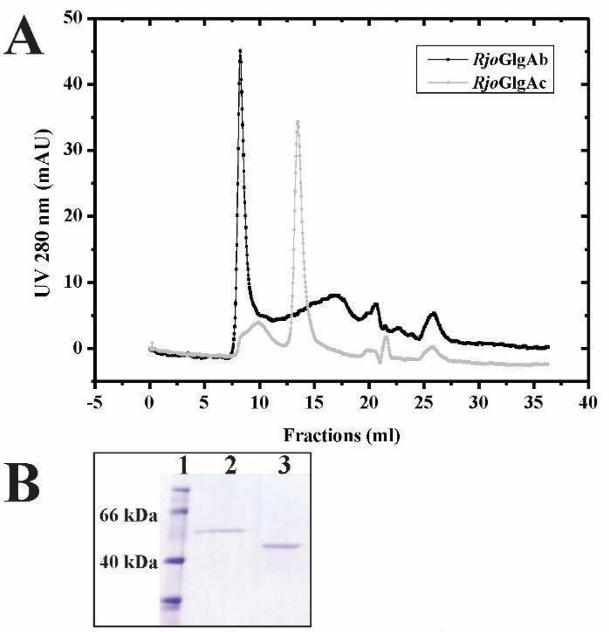


C

| | Activity (U/ml) |
|---|---|
| *Rjo*GlgAb | 0.021 |
| *Rjo*GlgAc | 0.027 |

**<u>Supplemental Figure 2</u>:** Gel filtration chromatogram (A) for both GlgAs. SDS-PAGE (B) for *Rjo*GlgAb (2) and *Rjo*GlgAc (3) peaks collected from the gel filtration analysis, and activity (C) for both enzyme peaks.

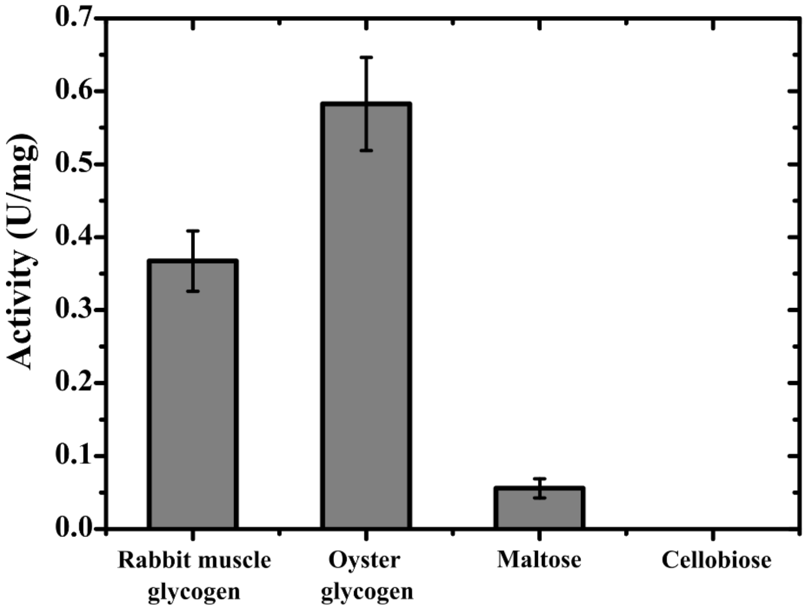


**<u>Supplemental Figure 3</u>:** Activity of *Rjo*GlgAb using different glucan molecules assayed: 2 mg/ml rabbit muscle glycogen, 2 mg/ml oyster glycogen type II, 2 mM maltose and 2 mM cellobiose, with 2 mM ADP-Glc.

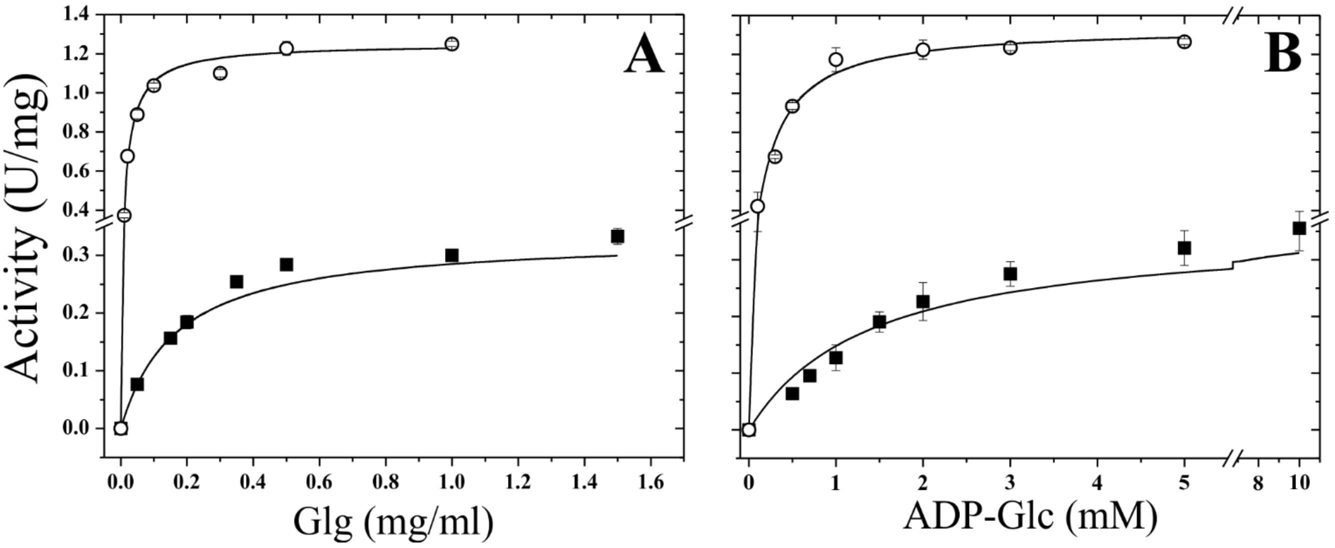


**<u>Supplemental Figure 4</u>:** *Rjo*GlgAb (filled squares) and *Rjo*GlgAc (open circles) saturation curves for glycogen (A) and ADP-Glc (B). A concentration of 1 mg/ml and 0.5 mg/ml of rabbit Glg were used for ADP-Glc saturation curves of *Rjo*GlgAb and *Rjo*GlgAc, respectively. For rabbit Glg saturation curves, 5 mM (*Rjo*GlgAb) and 1 mM (*Rjo*GlgAc) of ADP-Glc were used.

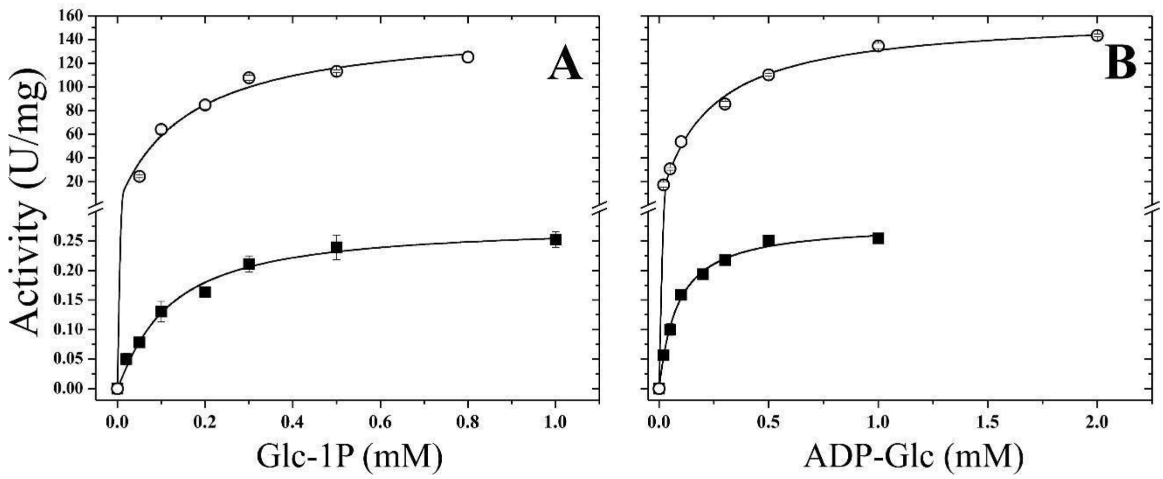


**<u>Supplemental Figure 5</u>:** Saturation curves for Mal-1P synthesis of *Rjo*GlgAb (filled squares) and *Rjo*GlgAc (open circles) with Glc-1P (A) and ADP-Glc (B). 1 mM of ADP-Glc was used for Glc-1P saturation curves for both enzymes, while the concentration of Glc-1P in both ADP-Glc saturation curves was 0.5 mM.
